# Inductive biases of neural specialization in spatial navigation

**DOI:** 10.1101/2022.12.07.519515

**Authors:** Ruiyi Zhang, Xaq Pitkow, Dora E Angelaki

## Abstract

The brain may have evolved a modular architecture for reward-based learning in daily tasks, with circuits featuring functionally specialized modules that match the task structure. We propose that this architecture enables better learning and generalization than architectures with less specialized modules. To test this hypothesis, we trained reinforcement learning agents with various neural architectures on a naturalistic navigation task. We found that the architecture that largely segregates computations of state representation, value, and action into specialized modules enables more efficient learning and better generalization. Behaviors of agents with this architecture also resemble macaque behaviors more closely. Investigating the latent state computations in these agents, we discovered that the learned state representation combines prediction and observation, weighted by their relative uncertainty, akin to a Kalman filter. These results shed light on the possible rationale for the brain’s modular specializations and suggest that artificial systems can use this insight from neuroscience to improve learning and generalization in natural tasks.

## Introduction

Accurate generalization beyond training tasks requires correct prior knowledge of the task structure [1]. However, as David Hume famously highlighted in his “problem of induction”, one’s prior knowledge can be fallacious, leading to unsuccessful generalization [2]. Nevertheless, animals have the ability to efficiently acquire the structure of their daily tasks as prior knowledge for novel tasks outside their typical domain [1, 3, 4, 5]. This remarkable ability may stem from the brain’s innate biases evolved for their daily tasks [1, 6, 7].

In theory, numerous solutions exist for a given task. For instance, one solution may involve comprehending the task’s underlying structure, whereas another could rely on memorizing all input-output pairs. Although both can produce positive results with sufficient training, a solution that understands the task structure is expected to exhibit greater data efficiency in mastering the training task and can generalize to new, structurally similar tasks; in contrast, rote memorization requires seeing all training examples and lacks generalizability. Every learning system for a task, whether biological or artificial, has a bias that favors some solutions over others, known as the inductive bias. For a neural network, its architecture defines a crucial aspect of this bias [6, 8, 9]. To prioritize solutions that learn the task structure and support generalization, the inductive bias must be tailored to the specific tasks of interest, as there is no universal inductive bias suitable for all tasks [10].

The remarkable ability of animals to rapidly learn and generalize in natural tasks suggests that their brains are indeed endowed with suitable inductive biases for these tasks. One perspective suggests that the brain may have evolved a modular architecture, allowing it to use functionally specialized computational modules that are appropriate for task requirements [11, 12, 13, 14, 15]. Each module specializes in a specific aspect of demanding computations, collectively covering all aspects of the task structure. We hypothesize that this architecture enables higher efficiency in learning the task structure, and better generalization, than those with less specialized modules.

To test this hypothesis, it is essential to select a natural task and compare a modular architecture designed for the task against alternative architectures. We chose a naturalistic virtual-navigation task previously used to investigate the neural computations underlying macaques’ flexible behaviors [16, 17]. Remarkably, macaques trained in this task demonstrated immediate generalization to novel tasks with a similar structure, without requiring any additional improvement [18]. In this task, subjects are required to steer towards a transiently visible target using optic flow cues. They must mentally compute multiple variables, including the internal state representation of the outside world (a ‘belief’) given partial and noisy sensory cues, the motor commands (actions) used to control a joystick for navigation based on the belief, and the value of each action [19].

In neuroscience, it would be hard to compare different architectures for solving a given task: we cannot easily control brain architecture, nor can we find multiple animals whose brains differ only in the computations of interest. However, we can explore behaviors of computational models with distinct architectures, since the principles governing the inductive bias of a learning system apply to both biological and artificial intelligence. Inspired by the brain’s modularity, we designed architectures where neurons are segregated into distinct modules, each dedicated to specific task variables. We also designed a set of other architectures for comparison, each featuring modules with weaker specializations for particular task variables.

To train artificial agents using these architectures, we used reinforcement learning (RL) [20] with sparse reward signals, similar to the training of macaques. Afterward, we evaluated these agents in two novel tasks that share a similar structure to the training task. One manipulated the sensorimotor mapping from joystick movements to subjects’ movements in the environment. The other randomly applied passive perturbations to subjects’ movements. We found that the architecture with highly specialized modules was more suitable for the navigation task than other architectures. This architecture efficiently established a more accurate understanding of the task structure, resulting in better and more animal-like behaviors in the training task. Moreover, this acquired knowledge facilitated a more accurate state representation in novel tasks, thus supporting instant generalization, similar to macaques’ ability in these tasks [18, 17].

Additionally, we found that the modular agent trained in this task exhibited a belief update rule resembling recursive Bayesian estimation [21]. Specifically, the posterior belief is calculated by weighing the prior prediction using a motor efference copy against a likelihood over states derived from visual observation. The reliability of the two sources is duly considered, with the more reliable source assigned a higher weight, akin to the brain’s probabilistic inference [22]. Consequently, agents trained with greater uncertainty in observation than in prediction tended to disregard observations in belief formation, which then impaired their generalization ability in tasks that required the use of novel observations to construct beliefs. Hence, similar to macaques and humans in this task [23], agents must develop prior knowledge of relying more on observation for generalization.

Together, these findings reinforce the ongoing endeavor to bridge the gap between artificial intelligence (AI) and neuroscience [6, 8, 24, 25]. From AI to neuroscience, we demonstrated how artificial models can be employed to explore hypotheses of the brain that are currently challenging to investigate through direct experimentation. From neuroscience to AI, we exemplified how drawing inspiration from the brain can empower artificial models to achieve better learning and generalization in natural tasks. This bi-directional exchange underscores the potential for mutually beneficial collaborations that can advance both fields.

## Results

### RL agents trained to navigate using partial and noisy sensory cues

To study naturalistic, continuous-time computations involved in foraging behaviors, we previously designed a virtual reality navigation task where macaques navigate to targets using sparse and transient visual cues [16]. At the beginning of each trial, the subject is situated at the origin facing forward; a target is presented at a random location within the field of view on the ground plane and disappears after 300 ms. The subject can freely control its linear and angular velocities with a joystick to move in the virtual environment (Fig. 1**a**). The objective is to navigate toward the memorized target location, then stop inside the reward zone—a circular region centered at the target location with a radius of 65 cm. A reward is given only if the subject stops inside the reward zone. The subject’s self-location is not directly observable because there are no stable landmarks; instead, the subject needs to use optic flow cues on the ground plane to perceive self-motion and perform path integration. Note that optic flow cues cannot serve as stable landmarks because ground plane textural elements—small triangles—appear at random locations and orientations, and disappear after only a short lifetime (∼ 250 ms). A new trial starts after the subject stops moving or the trial exceeds the maximum trial duration. Details of this task are described in [16]. Macaques can be trained to master this task, and all macaque data presented in this paper were adapted from previously published works [16, 17, 18, 23, 26, 27].

**Figure 1.**
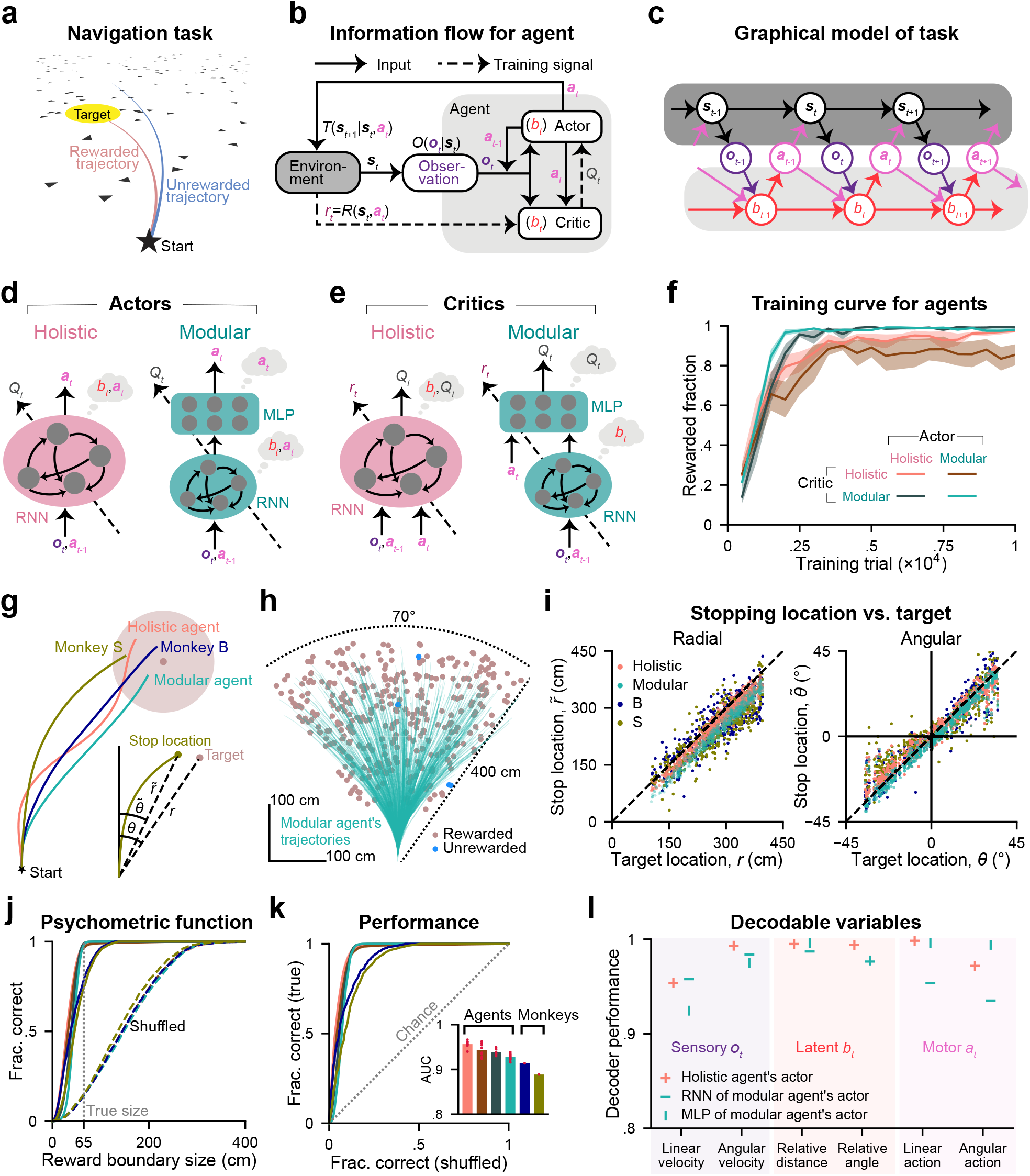
RL agents with different neural architectures were trained in a partially observable navigation task. **a**. Schematic of the navigation task from the subject’s perspective. Optic flow cues are generated by the motion of many randomly positioned and oriented triangle elements on the ground. These cues cannot serve as landmarks since each one exists for only a short time. A trial is rewarded only if the subject stops in the reward zone, and is otherwise unrewarded. **b**. Block diagram showing the interaction between an RL agent and the task environment. At each time step *t*, a partial and noisy observation ***o***_*t*_ of the state ***s***_*t*_ drawn from the distribution *O*, and the last action ***a***_*t*−1_ are provided to the actor and critic to form the belief *b*_*t*_ that is encoded in their network states ***h***_*t*_. The actor outputs an action ***a***_*t*_ based on *b*_*t*_, and this action updates ***s***_*t*_ to ***s***_*t*+1_; the critic estimates the value *Q*_*t*_ of this action based on *b*_*t*_. The actor is trained to generate the action that maximizes *Q*_*t*_; the critic updates its synaptic weights using the TD error after receiving reward *r*_*t*_ from the environment. **c**. Graphical model of the task. In the environment (dark gray), ***s***_*t*_ changes over time following the transition probability. Internally (light gray), *b*_*t*_ evolves following the belief update rule using observations ***o***_*t*_ and motor efference copies ***a***_*t*−1_. Observations ***o***_*t*_ are drawn given ***s***_*t*_. Actions ***a***_*t*_ are taken given *b*_*t*_. **d**. Schematic of actors with a holistic (*left*) or modular (*right*) architecture. A holistic actor uses an RNN to compute *b*_*t*_ and ***a***_*t*_ jointly; a modular actor consisting of an RNN and an MLP constrains the computation of *b*_*t*_ to the RNN, and leaves the MLP only to compute ***a***_*t*_. Thought bubbles denote the variables computed in each module. **e**. Similar to **d**, but for critic networks computing *b*_*t*_ and *Q*_*t*_. **f**. Fraction of rewarded trials during the training process following training phase I (see definition in Methods). The measurement is based on a validation set (500 trials) for each agent at each checkpoint, which occurs every 500 training trials. Shaded regions denote ± 1 SEM across training runs with *n* = 8 random seeds. **g**. An example trial showing monkeys and agents navigating toward the same target from the start location. Shaded circle: reward zone. Inset compares the target location (radial: *r*, angular: *θ*) vs. the stop location of monkey S (radial: 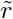, angular: 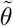). **h**. Overhead view of the spatial distribution of 500 representative targets and an example modular agent’s trajectories while navigating toward these targets. **i**. Comparison of agents/monkeys’ stop locations for the target locations from **h**. Black dashed lines have a slope of 1. **j**. Fraction of correct trials in a test set (1657 trials) as a function of hypothetical reward boundary size. Solid lines denote true data; dashed lines denote shuffled data. The gray dotted line denotes the true reward boundary size. **k**. True data versus shuffled data in **j** (ROC curve). Inset shows the area under the ROC curve (AUC). **j**–**k**. Agents’ data are averaged across *n* = 8 training runs. **l**. Performance of linear decoders trained to decode task variables from example neural modules using trials in **j**, with 70% for decoder training and the remaining for analyses. Performance is quantified by computing the Pearson’s *r* between true and decoded values of variables.

RL [20] is a reasonable framework for modeling behavior in this task because, like animals, RL agents can learn this task through sparse reward signals. We formulate this task as a Partially Observable Markov Decision Process (POMDP) [28] in discrete time, with continuous state and action spaces (Fig. 1**b**). At each time step *t*, the environment is in the state ***s***_*t*_ (including the agent’s position and velocity, and the target’s position). The agent takes an action ***a***_*t*_ (controlling its linear and angular velocities) to update ***s***_*t*_ to the next state ***s***_*t*+1_ following the environmental dynamics given by the transition probability *T* (***s***_*t*+1_ | ***s***_*t*_, ***a***_*t*_), and receives a reward *r*_*t*_ from the environment following the reward function *R*(***s***_*t*_, ***a***_*t*_) (a positive scalar if the agent stops inside the reward zone). The state ***s***_*t*_ is not fully observable, so to support decision-making, the agent needs to maintain an internal state representation that synthesizes the history of evidence. For example, a recursive Bayesian estimation (Methods) can form a posterior density *p*(***s***_*t*_ | ***o***_0:*t*_, ***a***_0:*t*−1_) over the current world state ***s***_*t*_ given the history of two sources of evidence. One source is a prediction based on its internal model of the dynamics, its previous posterior, and the last self-action ***a***_*t*−1_ (e.g., a motor efference copy). The other source is a partial and noisy observation ***o***_*t*_ of the state ***s***_*t*_ drawn from the observation probability *O*(***o***_*t*_ | ***s***_*t*_). For our task, this observation includes the target location when it is visible, and information about the agent’s movement estimated via optic flow. We call the resultant posterior the agent’s “belief” *b*_*t*_. We will show later that agents using neural networks trained in this task approximately encode this belief *b*_*t*_ in their neural activity ***h***_*t*_, and its dynamics are consistent with approximate recursive Bayesian estimation.

Here we use an *actor* -*critic* approach to learning [20] (Fig. 1**b**), where actor and critic are implemented using distinct neural networks trained end-to-end, and each network can individually learn to encode *b*_*t*_ in its neural activity (Methods). At each *t*, the actor computes the belief *b*_*t*_ of the state ***s***_*t*_ using two sources of evidence ***o***_*t*_ and ***a***_*t*−1_, and generates an action ***a***_*t*_. It is trained to generate the action that maximizes the state-action value *Q*_*t*_—the expected discounted cumulative rewards from *t* until the trial’s last step, given taking ***a***_*t*_ in ***s***_*t*_. Since the ground truth value is unknown, the critic computes the belief *b*_*t*_ of ***s***_*t*_ and estimates the value *Q*_*t*_, learned through the temporal-difference reward prediction error (TD error) after receiving the reward *r*_*t*_ (| *r*_*t*_ + *γQ*_*t*+1_ − *Q*_*t*_ |, *γ* denotes the discount factor, see Methods; Fig. 1**b**). The actor is updated more slowly than the critic, allowing the actor to consistently learn something new from the critic [29]. Over time, as the state evolves in the outside world following the environmental dynamics, the belief evolves in parallel following the internal belief update rule (learned through training). Sequential observations are drawn given the outside ***s***_*t*_; sequential actions are taken based on the internal *b*_*t*_ (Fig. 1**c**). Full details of this formulation are shown in Methods.

Actor and critic networks can use various architectures. Our objective here is to test the hypothesis that functionally specialized modules offer advantages for our task. Therefore, we designed architectures incorporating modules with distinct levels of specialization for comparison. We achieve this with two ways of confining certain variable computations to specific modules by restricting the memory or inputs of the modules. First, we confine the computation of beliefs for the actor and critic. Computing beliefs about slowly varying latent states requires synthesizing evidence over time, so a network that computes the belief must have some form of memory. Recurrent neural networks (RNNs) satisfy this requirement using a hidden state ***h***_*t*_ that evolves over time. In contrast, computations of value and action do not need memory when the belief is provided, making memoryless multi-layer perceptrons (MLPs) sufficient. Consequently, by adopting an architecture with an RNN followed by an MLP that cannot integrate signals over time, the computation of belief is exclusively confined to the RNN. Second, we confine the computation of the state-action value *Q*_*t*_ for the critic. By using a critic architecture with two sequentially connected modules, and by supplying the action input ***a***_*t*_ only to the second module, we restrict the computation of *Q*_*t*_ exclusively to that second module. This is because *Q*_*t*_ is necessarily a function of the action ***a***_*t*_, but the first module lacks access to the action so it cannot compute *Q*_*t*_.

Based on these ideas, we designed two types of architectures for both the actor and critic (more architectures will be shown later). The first is a *holistic* actor/critic, which comprises a single module where all neurons jointly compute the belief and the action/value. The second is a *modular* actor/critic, which confines the belief computation to an RNN module and dedicates an MLP module to action/value computations (Fig. 1**d**–**e**). Thought bubbles in Fig. 1**d**–**e** denote which variables can be computed within each module, determined by the aforementioned types of segregation. We trained agents using all four combinations of these two actor and critic types (Fig. 1**f**, legend). We refer to an agent whose actor and critic are both holistic or both modular as a holistic agent or a modular agent, respectively. The training concluded after agents had experienced 10^4^ trials (after training phase I, see Methods). This number of training trials was comparable to the experience of animals in this task and proved sufficient for the agents to achieve stable and accurate performance (Fig. 1**f**). Agents with modular critics demonstrated greater consistency across various random seeds (Fig. 1**f**, shaded regions) and achieved near-perfect accuracy more efficiently than agents with holistic critics.

Agents’ behavior was compared with that of two monkeys (Fig. 1**g**) for a representative set of targets uniformly sampled on the ground plane (distance: 100 to 400 cm, angle: − 35^*◦*^ to +35^*◦*^; modular/holistic agent: Fig. 1**h**/Fig. S1**a**). In the next section we will contrast the properties of agents’ trajectories, but first, we focus on the accuracy of their stop locations (linear: 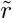, angular: 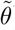) versus the target location (linear: *r*, angular: *θ*; Fig. 1**g**, inset). The tight correspondence between stop and target locations indicates that, similar to monkeys, all agents had learned the training task (Fig. 1**i**; Pearson’s *r*: Fig. S1**b**). When stop locations were regressed against target locations (without intercept), we noticed that, similar to monkeys, agents also systematically undershot targets (Fig. S1**c**: regression slope *<* 1). This finding can be predicted based on the RL framework: although the immediate reward for stopping at any location within the reward zone is the same, those considering long-term values discounted over time should prefer closer reward locations to save time.

We used a Receiver Operating Characteristic (ROC) analysis [16, 17] to systematically quantify behavioral performance. A psychometric curve for stopping accuracy is constructed from a large representative dataset by counting the fraction of rewarded trials as a function of a hypothetical reward boundary size (radius 65 cm is the true size; infinitely small/large reward boundary leads to no/all rewarded trials). A shuffled curve is constructed similarly after shuffling targets across trials (Fig. 1**j**). Then, an ROC curve is obtained by plotting the psychometric curve against the shuffled curve (Fig. 1**k**). An ROC curve with a slope of 1 denotes a chance level (true=shuffled) with the area under the curve (AUC) equal to 0.5. High AUC values indicate that all agents reached good performance after training (Fig. 1**k**, inset). This performance can be explained by accurate task variables encoded in their neural networks (actor: Fig. 1**l**, critic: Fig. S1**d**; see Methods), as previously also shown in the macaque brain [17, 26, 27]. In summary, all agents with different neural architectures can be trained to perform our navigation task (Fig. 1**k**).

### Different architectures yield different performance in the training task

We have shown that all trained agents achieved high accuracy by analyzing their stop locations. However, we have also noticed distinct characteristics in trajectories across different agents (Fig. 1**g**). Here, we examined two crucial trajectory properties: curvature and length. When tested on the same series of targets as the monkeys experienced, agents with modular critics displayed more efficient trajectories than those with holistic critics, characterized by smaller curvature and length (Fig. S2**a**–**b**). Notably, the difference between trajectories generated by agents with modular critics and those of monkey B was comparable to the variation between trajectories of two monkeys (Fig. 2**a**–**b**). In contrast, when agents used holistic critics, the difference in trajectories from monkey B was much larger, suggesting that modular critics facilitated more natural behaviors.

**Figure 2.**
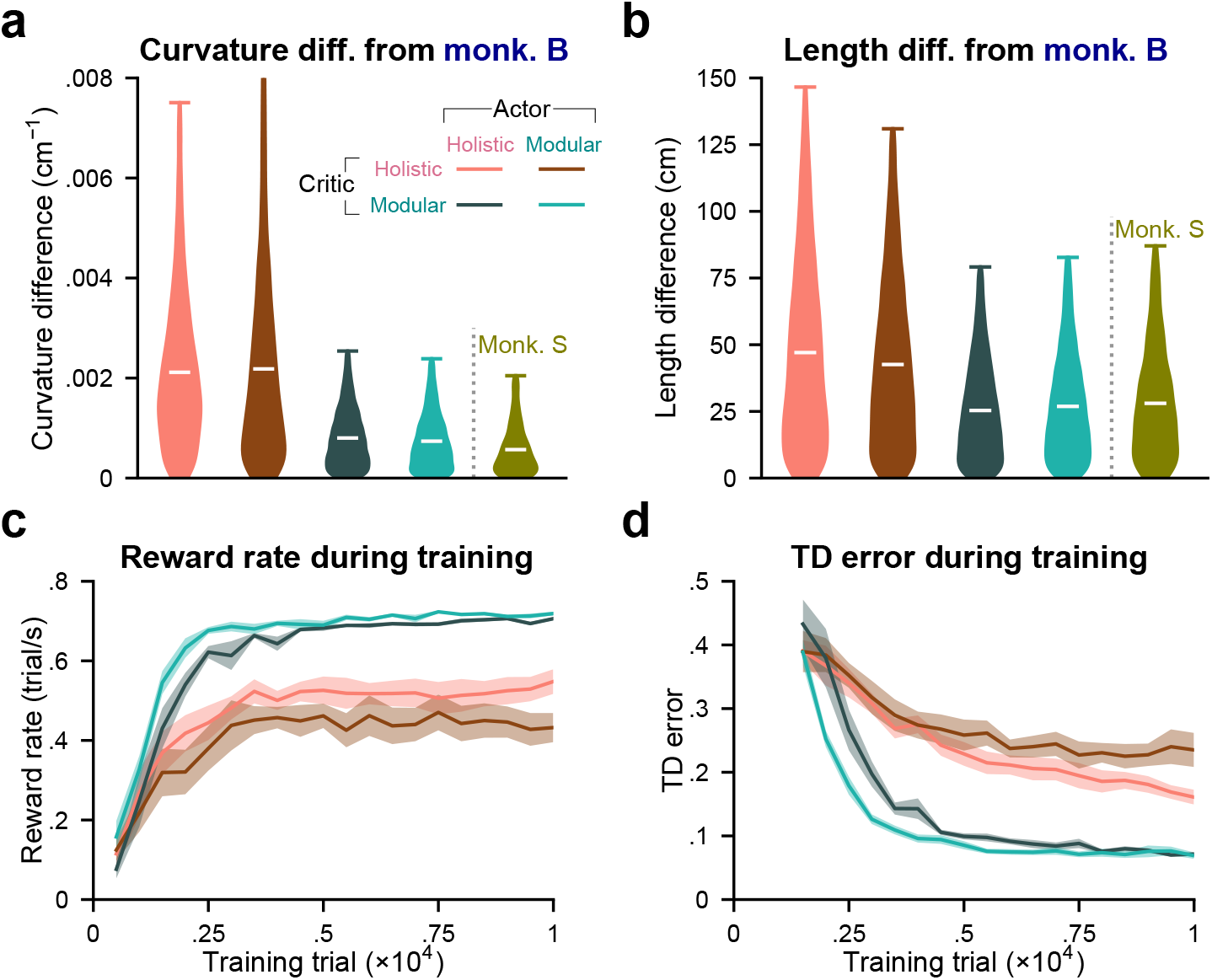
Agents with modular critics exhibit superior efficiency and performance in learning. **a**. Distribution of the absolute curvature difference between agents’ and monkey B’s rewarded trajectories, as well as the difference between two monkeys’ rewarded trajectories, while all navigated to the same set of 1000 targets. The curvature values were averaged across timesteps for each trajectory. **b**. Similar to **a**, but showing the absolute length difference for each trajectory. **a**–**b**. Containing data from *n* = 8 random seeds for each agent. White bars denote means across trials. **c**. Similar to Fig. 1**f**, but showing the reward rate. For each checkpoint, the reward rate is calculated by dividing the number of rewarded trials in a validation set (500 trials) by the time spent in seconds. **d**. Similar to **c**, but showing the agents’ TD error averaged across timesteps and trials in the validation set after they reached an average accuracy of 60% across seeds. At each step *t*, the critic computes *Q*_*t*_ based on the state and action at *t*, and *Q*_*t*+1_ based on the state and action at *t* + 1. The TD error is then |*r*_*t*_ + *γQ*_*t*+1_ − *Q*_*t*_|, where *r*_*t*_ and *γ* denote the current reward and the discount factor (Methods).

Agents are expected to develop efficient behaviors, as the value of their actions gets discounted over time. Therefore, we assess their efficiency throughout the training process by measuring the reward rate, which refers to the number of rewarded trials per second. We found that agents with modular critics achieved significantly higher reward rates (Fig. 2**c**), which explains their more efficient trajectories (Fig. S2**a**–**b**). Later, we will show that even with a training period ten times longer, agents with holistic critics still fail to match the reward rate attained by agents with modular critics within this short training period.

Since the actors responsible for generating actions were trained by maximizing the critics’ value estimation instead of the latent ground-truth value, the lower reward rates may be attributed to inaccurate value estimation. To investigate this, we monitored the TD error for critics during training. This error is the discrepancy between the current value estimate and the discounted subsequent value estimate combined with the current reward, serving as the learning objective for the critic ([20]; Methods; Fig. 2**d**). A critic that perfectly comprehends the structure of the task dynamics and rewards should yield no errors. Interestingly, agents with modular critics exhibited faster convergence of TD errors, ultimately reaching significantly lower values compared to agents with holistic critics (Fig. 2**d**). This suggests that the modular critic enhances efficiency and accuracy in learning the task structure, thereby providing a training signal that closely aligns with the true nature of the task for the actor. Consequently, actors with either architecture can develop superior behavior in the training task, as opposed to those trained by the holistic critic (Fig. 2**c**). Next, we will evaluate the generalization capabilities of these trained agents in novel tasks.

### Gain task: generalization to novel sensorimotor mappings

One crucial aspect of sensorimotor mapping is the joystick gain, which linearly maps motor actions on the joystick (dimensionless, bounded in [− 1, 1]) to corresponding velocities in the environment. During training, the gain remains fixed at 200 cm/s and 90^*◦*^/s for linear and angular components, referred to as the × 1 gain. By increasing the gain to values that were not previously experienced, we create a “gain task” manipulation. Remarkably, monkeys demonstrated immediate generalization to novel gains and other task manipulations without requiring further improvement [18]. This prompts us to investigate whether our trained agents can demonstrate similar generalization abilities.

To assess this, monkeys and agents (Methods, agent selection) were tested with novel gains of 1.5*×* and 2*×* (Fig. 3**a**). Blindly following the same action sequence as in the training task would cause the agents to overshoot (no-generalization hypothesis: Fig. 3**b**, dashed lines; Methods). Instead, the agents displayed varying degrees of adaptive behavior (Fig. 3**b**, solid lines). To quantitatively evaluate behavioral accuracy while also considering over-/under-shooting effects, we defined radial error as the Euclidean distance between the stop and target locations in each trial, with positive/negative sign denoting over-/under-shooting (using idealized trajectories, see Methods). Under the novel gains, agents with modular critics consistently exhibited smaller radial errors than agents with holistic critics (Fig. 3**c**), with the modular agent demonstrating the smallest errors, comparable to those observed in monkeys (Fig. 3**d**, Fig. S3**a**). ROC analyses further confirmed the performance differences among agents (Fig. 3**e**–**f**).

**Figure 3.**
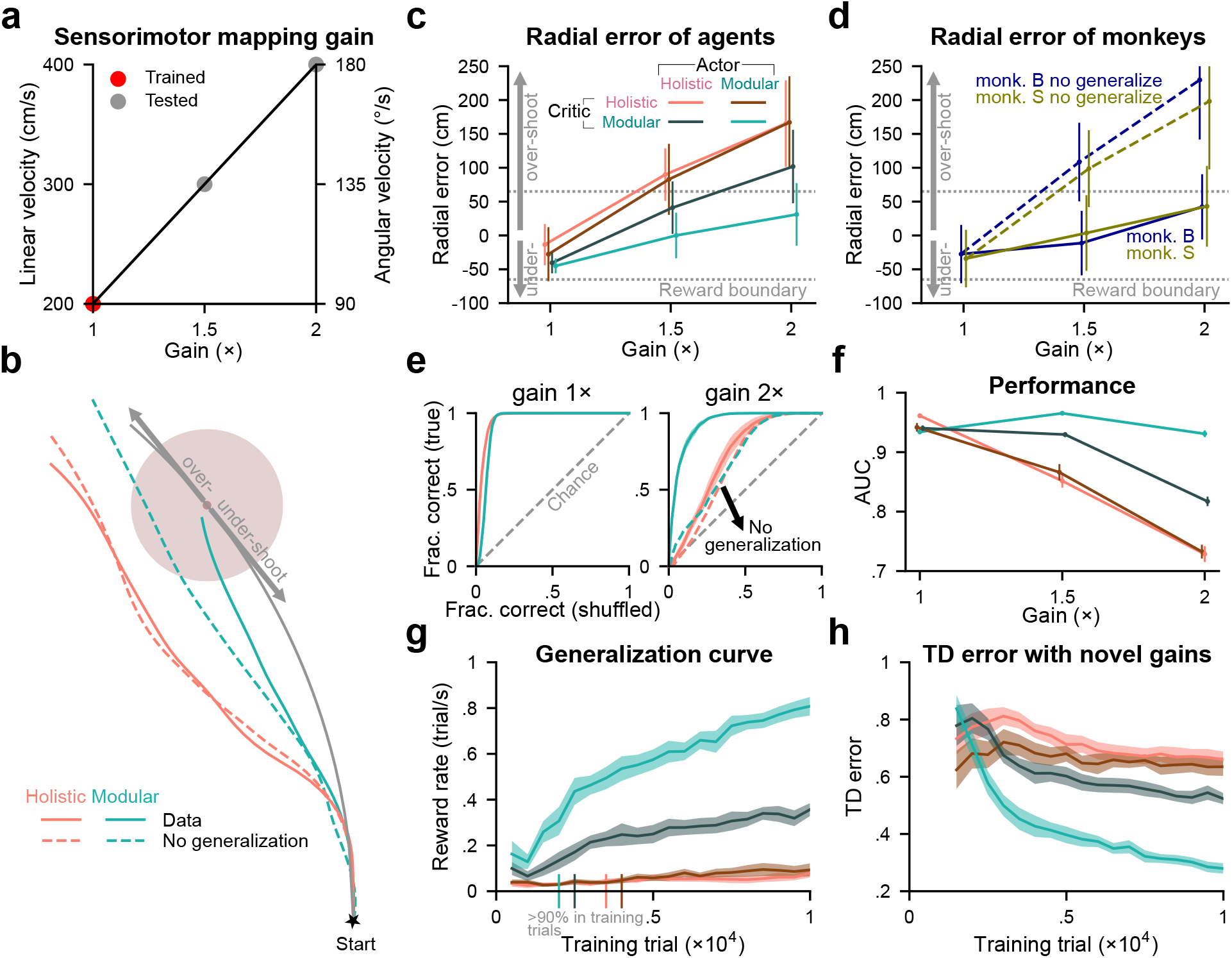
The modular agent exhibits the best generalization performance in the gain task. **a**. Gain is the parameter that linearly maps the joystick actions (bounded in [−1, 1]) onto velocities in the environment. 1× gain used in training has linear and angular components of 200 cm/s and 90 ^*◦*^/s, respectively. After training, animals and agents were tested with 1.5× and 2× gain values. **b**. Example trajectories of agents navigating toward a target with a novel 1.5× gain. Dashed lines denote hypothetical no-generalization trajectories (Methods). Arrows indicate regions of over- or under-shooting relative to the distance along an idealized circular trajectory connecting the start location to the target (grey line). **c**. Radial error of agents’ stop locations across different gains. The absolute value of a trial’s radial error is defined as the distance between the subject’s stop and target locations. The positive/negative sign of the radial error denotes over-/under-shooting (Methods). For each gain and agent, 8000 trials were conducted by concatenating results from *n* = 8 random seeds, with the same set of 1000 targets for each seed. Error bars denote ±1 SD across trials. The gray dotted lines denote the reward boundary. **d**. Similar to **c**, but for monkeys (the same 1000 targets as in **c** for each gain and monkey). **e**. ROC curves quantifying the accuracy of agents’ stop locations in different gain conditions. Gray dashed lines denote the chance level; dashed lines in other colors denote agents’ hypothetical no-generalization hypotheses. **f**. Area under the ROC curves in **e** (AUC) for all agents and gain conditions. **g**–**h**. Similar to Fig. 2**c**–**d**, but data were obtained by averaging over two validation sets (the same set of 500 targets, gain= 1.5×, 2×). Vertical bars overlaid on the x-axis in **g** denote the first time agents reached 90% accuracy in the 1× validation set (the same 500 targets, averaged across seeds). **e**–**h**: Lines denote means across *n* = 8 random seeds for each agent; shaded regions or error bars denote *±*1 SEM.

Similar to the assessment of the agents’ trajectory characteristics, reward rates, and TD errors in the previous section (Fig. 2), we again evaluated these quantities, this time under the novel gains. Agents with modular critics displayed more animal-like generalization trajectories, higher reward rates, and lower TD errors than agents with holistic critics, with the modular agent showing the closest resemblance to monkeys, the highest reward rates, and the lowest TD errors (Fig. S3**b**–**c**, Fig. 3**g**–**h**). Notably, the modular agent not only learned the training task the fastest (Fig. 3**g**, vertical bars on the x-axis) but also learned to generalize better and faster than other agents, continuing to improve its generalization with additional training trials (Fig. 3**g**). This trend is also evident in the accuracy of the agents’ value estimates on novel gain trials (Fig. 3**h**).

Together, these results demonstrate the impact of different inductive biases on generalization to novel gains. The modular critic enables better generalization than the holistic critic, and the combination of a modular critic and modular actor produces the best generalization performance.

### Accuracy of agents’ internal beliefs explains generalization in the gain task

Although we have confirmed that agents with distinct neural architectures exhibit varying levels of generalization in the gain task, the underlying mechanism remains unclear. We hypothesized that agents with superior generalization abilities should generate actions based on more accurate internal beliefs within their actor networks. Therefore, the goal of this section is to quantify the accuracy of beliefs across agents tested on novel gains, and to examine the impact of this accuracy on their generalization performance.

During the gain task, we recorded the activities of RNN neurons in the agents’ actors, as these neurons are responsible for computing the beliefs that underlie actions (Fig. 1**d**). As expected, these neurons showed sensitivity to the agents’ locations within the environment (spatial tuning; holistic agent: Fig. 4**a**, modular agent: Fig. 4**b**; Methods). To systematically quantify the accuracy of these beliefs, we used linear regression (with *ℓ*_2_ regularization) to decode agents’ locations from the recorded RNN activities for each gain condition (Fig. 4**c**; Methods). We defined the decoding error, which represents the Euclidean distance between the true and decoded locations, as an indicator of belief accuracy. While all agents demonstrated small decoding errors under the training gain, we found that agents struggling with generalization under increased gains (Fig. 3**f**) also displayed reduced accuracy in determining their own location (Fig. 4**d**, Fig. S4**a**). In fact, agents’ behavioral performance correlates with their belief accuracy (trial-average: Fig. 4**e**; trial-by-trial: Fig. S4**b**), a trend that was also observed in monkeys [17].

**Figure 4.**
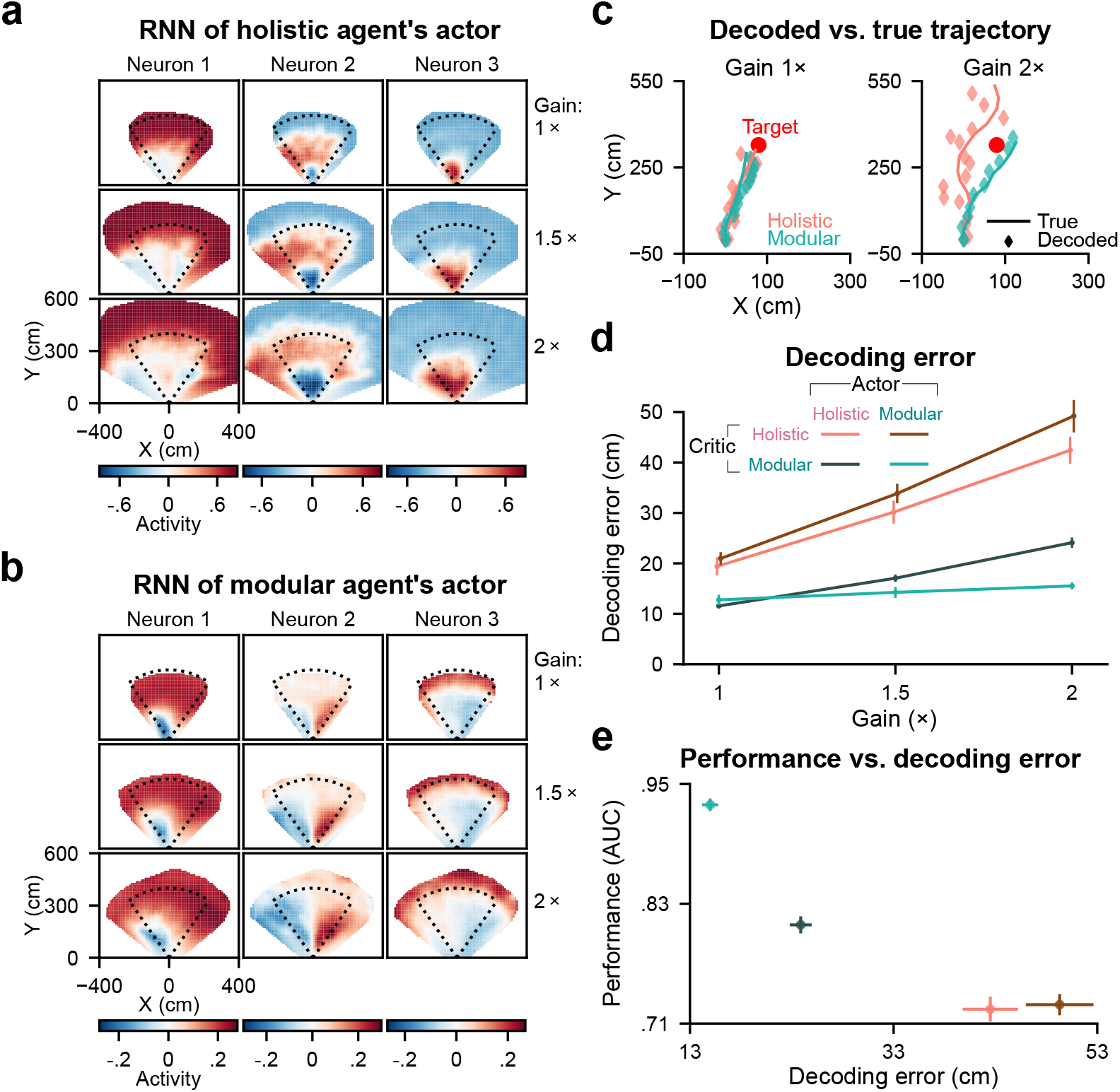
Decoding error of agents’ internal beliefs correlates with their behavioral performance in the gain task. **a**. Spatial tuning of example RNN neurons in a holistic agent’s actor. Each column denotes a neuron; each row denotes a gain condition (2000 trials). Dotted lines denote the boundary of a region containing all target locations. **b**. Similar to **a**, but for example RNN neurons in a modular agent’s actor. **c**. Decoded belief trajectories versus agents’ true trajectories during navigation to an example target under different gain conditions. Agents’ belief trajectories were estimated by linear decoders trained to decode agents’ locations from RNNs’ activities in actors (Methods). **d**. Decoding error as a function of gain. This error is defined as the distance between the true and decoded locations at each time step, and is averaged across time steps and trials in the test set. **c**–**d**. For each seed of each agent in each gain condition, 1400 trials were used to train a decoder, and 600 new trials were used for analyses. **e**. AUC versus decoding error for 2× gain. For each seed of each agent, 3500 trials were used to train a decoder, and 1500 new trials were used for analyses. **d**–**e**: Error bars denote *±*1 SEM across *n* = 8 random seeds.

These analyses suggest that the inductive biases of neural architectures impact generalization performance by determining the accuracy of agents’ internal state representation in the novel gain task after being trained in the training task. Among all agents, the modular agent generalizes the best, supported by its most accurate internal belief.

### Perturbation task: generalization to passive motions, explained by belief accuracy

To assess one’s ability for generalization with manipulated latent states in the environment, we introduce another novel task called the “perturbation task” (Methods). This task involves randomly applying passive perturbation velocities to the joystick control for both the linear and angular components, causing monkeys or trained agents (Methods, agent selection) to deviate from their intended trajectories. The perturbations follow a Gaussian temporal profile lasting for one second, with parameters including peak time and peak magnitudes for the passive linear and angular velocities randomly sampled for each trial (Fig. 5**a**). Fig. 5**b** illustrates an example trial, displaying agents’ adaptive behaviors (solid: data, dashed: no generalization hypothesis, see Methods) in response to the sampled perturbations (inset).

**Figure 5.**
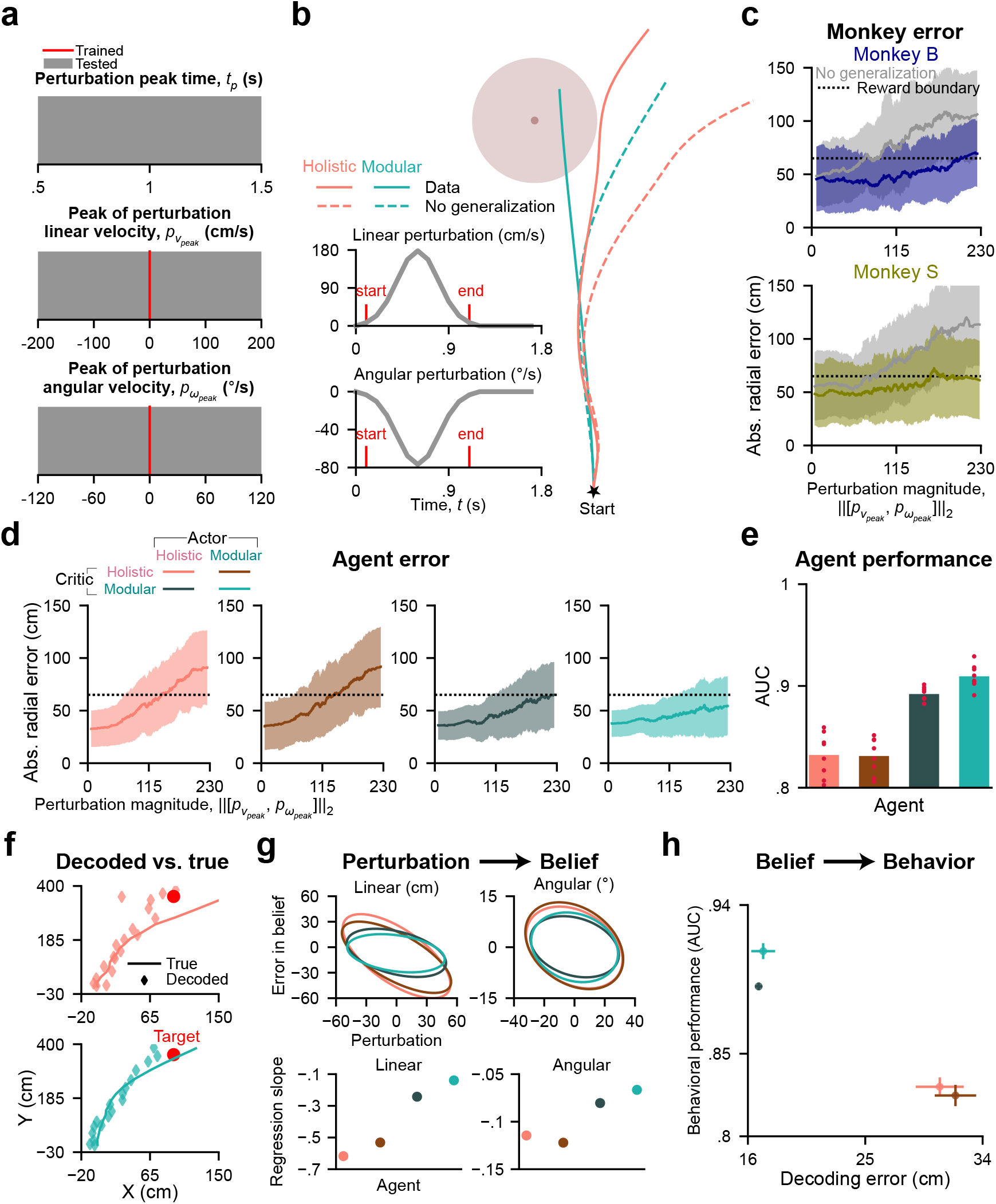
Agents with modular critics exhibit better generalization performance in the perturbation task, supported by their more accurate internal beliefs. **a**. A perturbation trial adds Gaussian-shaped passive velocities to joystick control at a random time for one second. In each trial, the perturbation peak time relative to the trial start (*top*) and the peak amplitudes of linear (*middle*) and angular (*bottom*) perturbation velocities are uniformly sampled from the corresponding ranges. No perturbations were used during training. **b**. Agents navigate in an example perturbation trial. Dashed lines denote hypothetical no-generalization trajectories (Methods). Inset shows linear (*top*) and angular (*bottom*) perturbations in this trial. **c**. Monkeys’ absolute radial error as a function of perturbation magnitude (the Euclidean norm of linear and angular perturbations) in 1000 trials. Errors for hypothetical no-generalization trajectories are in gray. Solid lines and shaded regions denote means and ± 1 SD obtained using a moving window (size=100 trials). The dotted black line denotes the reward boundary. **d**. Similar to **c**, but for artificial agents (1000 trials with target locations and perturbation parameters identical to those in **c** for each seed of each agent, resulting in 8000 trials for each agent; moving window size=800 trials). **e**. AUC for data with perturbation magnitude greater than 115 in **d**. Bars denote means across random seeds for each agent; red dots denote data for individual seeds. **f**. Similar to Fig. 4**c**, but showing trajectories navigating to an example target under perturbation. **g**. *Top*: Confidence ellipses of the deviation of decoded stop locations from true stop locations (*left*/*right*: using locations’ linear distance/polar angle) versus the integral of perturbation velocities (*left*/*right*: linear/angular perturbation) over trials. Data were produced by agents with 8 random seeds each and are shown in Fig. S5**e**–**f**. Confidence ellipses represent 1 SD of the data. Pearson’s *r* for agents’ data (in the same order as in **d**) for the left: − 0.67, − 0.69, − 0.42, − 0.30; for the right: − 0.30, − 0.31, − 0.26, − 0.19. *p* = 0. *Bottom*: Regression slope (without intercept) for the data. **h**. Similar to Fig. 4**e**, but for perturbation trials. **f** –**h**. For each seed of each agent, 3500 trials were used to train a decoder, and 1500 new trials were used for analyses. The peak magnitudes were sampled from the uniform distributions in **a**, but for **h**, smaller linear (within [−100, 100]) and angular ([−60, 60]) perturbations were excluded.

Monkeys displayed adaptation to perturbations, as evidenced by their behavioral errors (Fig. 5**c**). When faced with the same perturbations, agents with modular critics displayed errors comparable to those of monkeys, which were significantly smaller than the errors produced by agents with holistic critics (Fig. 5**d**). ROC analysis also supports these findings (Fig. 5**e**). This performance difference can be attributed to the agents’ ability to adjust behaviors: agents with modular critics demonstrated greater compensation for perturbations compared to those with holistic critics (Fig. S5**a**–**b**). Macaques and humans also exhibited compensatory behaviors in this task [23].

Similar to our observations in the gain task (Fig. 3**g**–**h**), agents’ generalization abilities (measured by reward rates) under perturbation improved with increased exposure to training trials (Fig. S5**c**), and those with higher reward rates demonstrated a better understanding of the perturbation task, as indicated by their lower TD errors (Fig. S5**d**).

We further investigated the neural mechanisms underlying agents’ different generalization abilities. As agents’ locations were perturbed, their internal beliefs should continuously track these perturbed locations. Failure to do so would introduce errors into their internal beliefs, ultimately impacting generalization behaviors. To test this, similar to our approach in the gain task (Fig. 4), we recorded the agents’ locations and the activities of RNN neurons in their actors under perturbations. We then linearly decoded the agents’ locations from these activities (Fig. 5**f** ; see Methods) and measured the difference between the true and decoded locations as an indicator of belief accuracy. We found that increased perturbations caused agents’ beliefs to deviate from the true locations (Fig. 5**g**, top; Fig. S5**e**–**f**), with agents using modular critics being less affected than those using holistic critics (Fig. 5**g**, bottom). These belief errors, akin to what we observed in the gain task (Fig. 4**e**, Fig. S4**b**), then propagated to behavioral errors (trial-average: Fig. 5**h**; trial-by-trial: Fig. S5**g**).

These analyses again demonstrate the impact of architectural inductive biases on generalization. By enabling agents to learn internal beliefs that remain accurate under novel perturbations, modular critics facilitate superior generalization than holistic critics.

### More architectures using less specialized modules

Above, we compared the holistic architecture against the modular architecture to investigate the advantages of module specialization. In this section, we aim to further strengthen our argument by introducing more architectures that deviate from the modular architecture (Fig. 6**a**–**b**).

**Figure 6.**
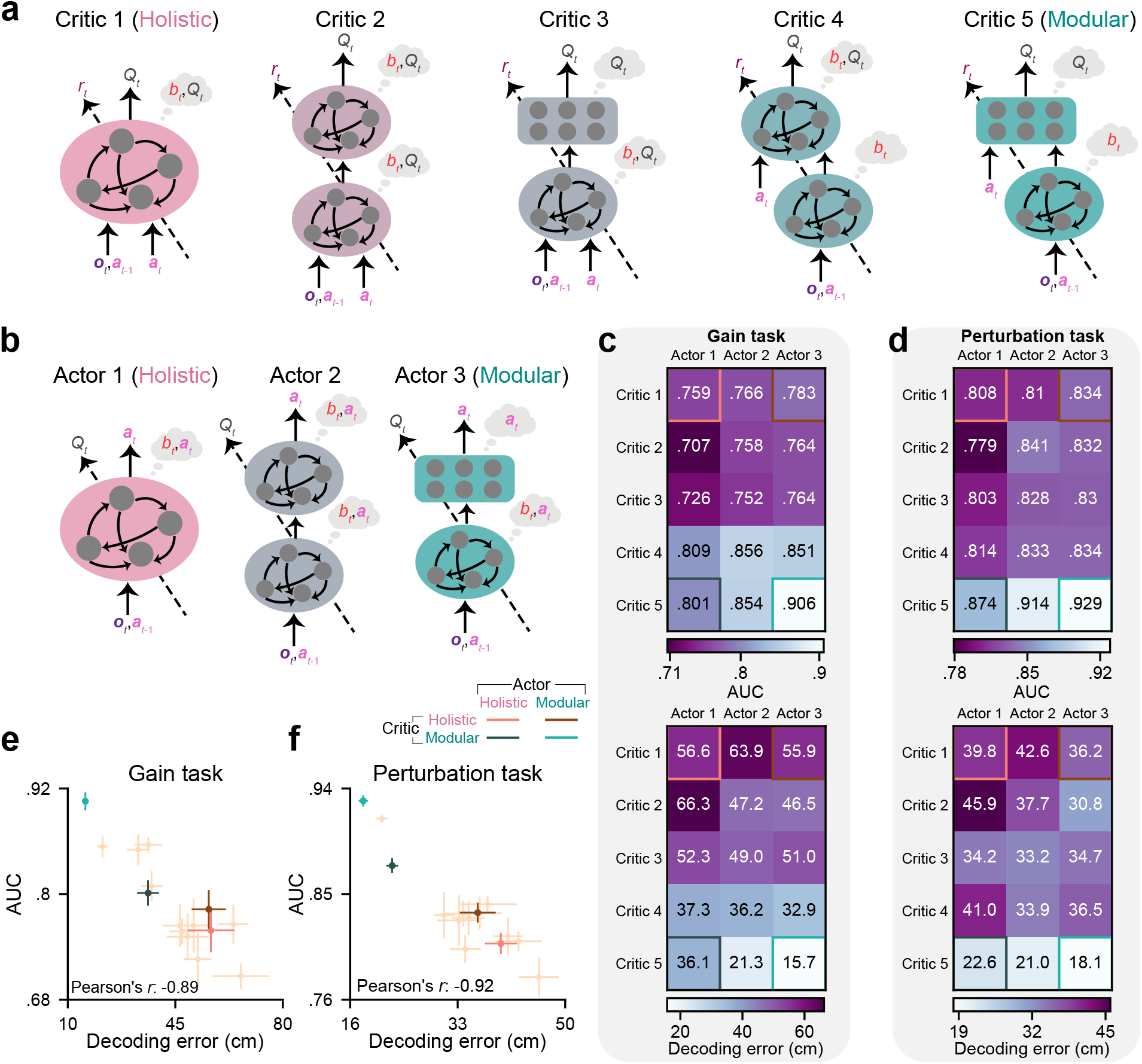
Agents using less specialized modules exhibit less accurate internal beliefs, resulting in inferior performance compared to the modular agent. **a**–**b**. Critic and actor diagrams as in Fig. 1**e** and 1**d**, but including more architectures with less specialized modules. **c**–**d**. AUC (*top*) and decoding error (*bottom*, averaged across time steps and trials) of agents in the gain (**c**) and perturbation (**d**) tasks, averaged across *n* = 8 random seeds. For each seed of each agent, 2000 trials were conducted. Gains were sampled from [3 ×, 4 ×] for **c**. Linear and angular perturbation velocities were sampled from [−200, 800], [−180, 180] for **d**. Beliefs were decoded from the RNN for actors 1 and 3, or the first RNN for actor 2. The four corners represent the four agents used in the previous analyses. Text in white/black denotes that the agent is worse/better than the average value of all agents. **e**–**f**. AUC versus decoding error using data in **c**–**d**. Error bars denote *±*1 SEM across random seeds.

By using two sequential RNNs instead of one, critic/actor 2 can distribute computations of two variables across two modules without enforced specialization. Substituting an MLP for the second RNN in critic/actor 2 yields critic/actor 3, where the belief computation over time is confined to the first RNN. Alternatively, we can retain two RNNs in the critic and exclusively provide the action input to the second RNN. This yields critic 4, where the value computation is confined to the second module. The modular critic confines both the belief and value computation and has the most module specialization. Note that the total number of trainable parameters is designed to be similar across all architectures (Fig. S6**a**, see Methods).

We trained agents using all combinations of these critics and actors, extending the training duration to ten times the previously used duration (10^5^ trials after training phase I, Methods). This allowed us to explore whether agents with inferior architectures could achieve comparable performance as those with good architectures through extensive training. We found that agents with less specialized critics still demonstrated lower reward rates than those with a modular critic (Fig. S6**b**, left). This can be attributed to the less accurate value estimates provided by their critics for training their actors (TD errors, Fig. S6**b**, right, Pearson’s *r* = −0.93).

We then compared these agents’ (Methods, agent selection) generalization abilities by conducting tests with challenging gain and perturbation parameters to obtain a robust evaluation and uncover nuances (Fig. 6**c**–**d**). Across critics, the modular critic (critic 5) outperformed all others. Note that the architectural components used in the pairs of critics 5 and 3 (RNN+MLP) and critics 4 and 2 (RNN+RNN) are identical. The only difference is that in critics 5 and 4, the value computation is confined to the second module, as only there can the computation access the action ***a***_*t*_, whereas critics 3 and 2 allow both modules to access the action, eliminating such confinement. The performance difference within each pair highlights the advantages of specialization for value computation. Similarly, in critic 5, the belief computation is confined to the first module, while in critic 4 it is not confined. The performance difference between critics 5 and 4 demonstrates that specialization for belief computation could further enhance performance. However, specializing in belief without specializing in value did not provide benefits, as indicated by the difference between critics 3 and 2 (AUC: Fig. 6**c**–**d**, top; reward rate: Fig. S6**c**–**d**).

Actors’ performance was contingent on the choice of critic architectures, as actors were trained by critics to learn the task structure. With a modular critic, the modular actor benefits from a dedicated belief module, enabling the development of an internal belief that remains accurate in novel tasks (Fig. 6**c**–**d**, bottom). Consequently, it outperformed others with less accurate beliefs (Fig. 6**e**–**f**).

Together, we conclude that the architecture using the modular critic and modular actor represents the most appropriate inductive bias for our task, benefiting learning and generalization.

### RNNs learn a Kalman filter-like belief update rule

We have thus far demonstrated that the modular agent has the most appropriate neural architecture to construct an accurate internal belief for our navigation task. Here, we explore this agent’s belief update rule using two information sources with uncertainties: observation (optic flow) and prediction (using a motor efference copy).

The Kalman filter [21] is a practical method to implement recursive Bayesian estimation when all variables are Gaussian, and the state transitions and observations are linear (Fig. 7**a**). It constructs an internal belief of states (posterior) using motor predictions (prior) and visual cues (likelihood). A prior predicts the state using the last self-action ***a***_*t*−1_ with uncertainty ***σ***_*a*_, the last belief *b*_*t*−1_ with uncertainty *P*_*t*−1_, and the state transition *T*. The posterior (belief *b*_*t*_) is a weighted average of the prior and the likelihood over states, given the observation ***o***_*t*_ with uncertainty ***σ***_*o*_. The weight is known as the Kalman gain, which weighs more on the source with a smaller uncertainty. In combining these sources of evidence, the posterior has an uncertainty *P*_*t*_ smaller than only relying on a single source. Note that for our task, the Kalman gain is only affected by ***σ***_*a*_ and ***σ***_*o*_, and is independent of the prior uncertainty *P*_*t*−1_ in prediction (see Methods).

**Figure 7.**
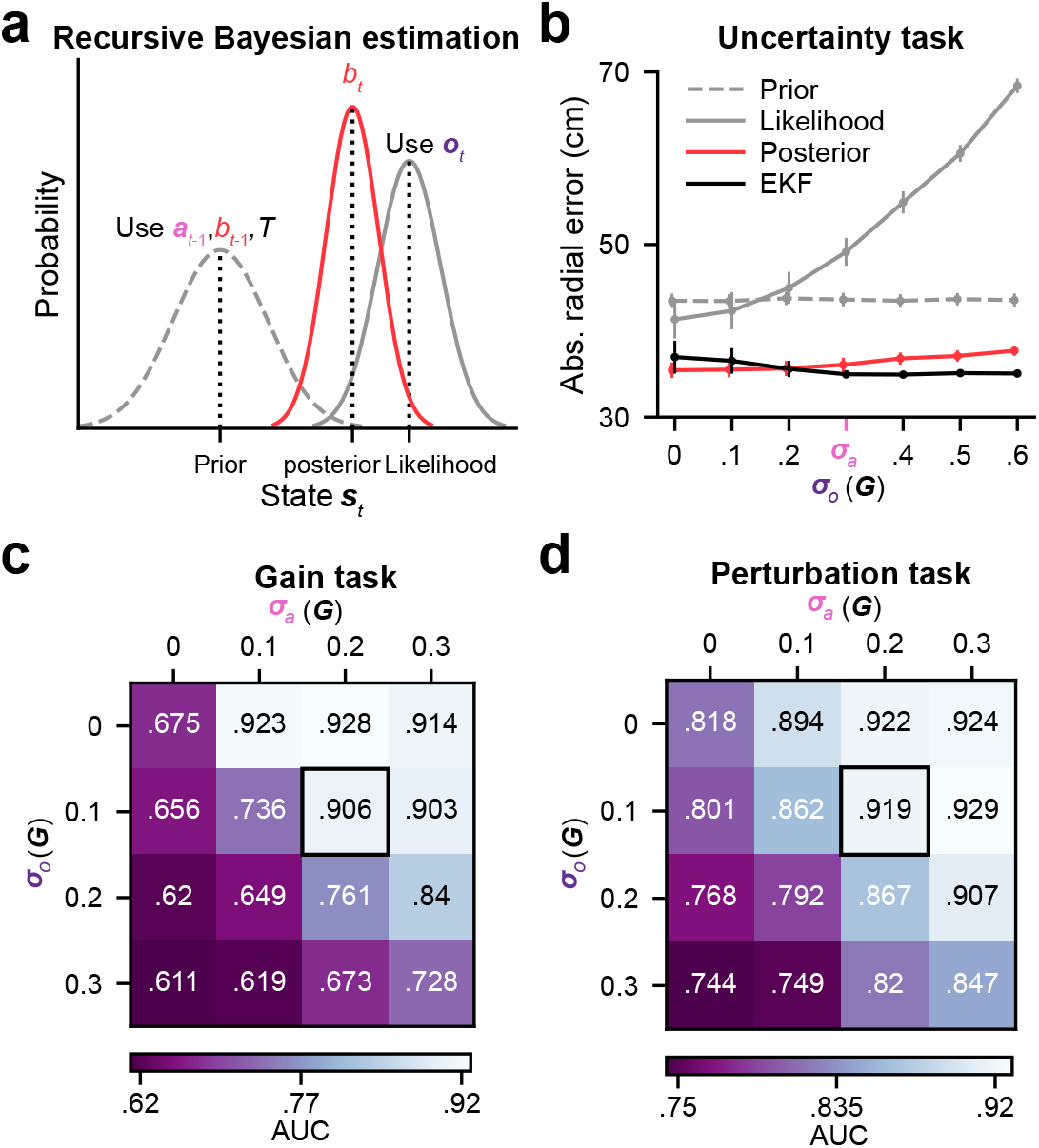
Modular agents with a Kalman filter-like belief update rule generalize worse in the gain and perturbation tasks when relying less on observation. **a**. Schematic of recursive Bayesian estimation (as implemented by the Kalman filter, Methods) in the 1D case, given zero-mean Gaussian noises. The prior is a prediction of the state ***s***_*t*_ using the last action ***a***_*t*−1_, the last belief *b*_*t*−1_, and the state transition *T*. The state’s likelihood depends on visual observation ***o***_*t*_. The posterior *b*_*t*_ combines these two sources and provides a state estimation with an uncertainty smaller than only relying on a single source. **b**. Absolute radial error of agents (*n* = 8 random seeds each) tested in the uncertainty task as a function of the observation uncertainty ***σ***_*o*_, given a fixed uncertainty in prediction ***σ***_*a*_. The prior model was trained with ***σ***_*a*_ = **0** ≪ ***σ***_*o*_. The likelihood model was trained with ***σ***_*o*_ = **0** ≪ ***σ***_*a*_. The posterior model was trained with ***σ***_*o*_ values that can be larger or smaller than ***σ***_*a*_ on each trial (see Methods and text). Error bars denote ±1 SEM across random seeds. **c**–**d**. AUC of agents trained with combinations of ***σ***_*a*_ and ***σ***_*o*_ tested in the gain (**c**) and perturbation (**d**) tasks, averaged across *n* = 8 random seeds for each agent. Gain and perturbation parameters were sampled uniformly from the same ranges as in Fig. 6**c**–**d**, but smaller linear (within [− 100, 100]) and angular ([− 60, 60]) perturbation velocities were excluded for **d**. Black boxes denote the combination used in the previous analyses as the default setting. **b**–**d**. 2000 trials were used for each seed of each agent for each uncertainty condition.

The RNN in an ideal modular agent may learn a belief update rule similar to the Kalman filter (more precisely, an extended Kalman filter [EKF] [30] allowing nonlinear state transitions; see Methods). We therefore compare modular agents with an EKF agent whose architecture is designed by replacing the modular actor’s and modular critic’s RNNs with an EKF. We trained and tested these agents in an “uncertainty task” (see Methods and below) to probe RNNs’ belief update rule. Uncertainties are represented in units of the joystick gain ***G***, and an uncertainty of **0** denotes the noise-free case.

We trained three types of modular agents under three uncertainty conditions. The first type is referred to as the prior model, trained with ***σ***_*a*_ = **0 ≪ *σ***_*o*_, which learns to only rely on prediction rather than the uninformative observation; the second type is the likelihood model, trained with ***σ***_*o*_ = **0 ≪ *σ***_*a*_, which learns to only rely on its perfectly accurate observation; the last one, the posterior model, is trained with a sampled value of ***σ***_*o*_ on each trial which can be larger or smaller than ***σ***_*a*_, and must learn to rely on both sources. Testing these models with ***σ***_*a*_ = 0.3***G*** and ***σ***_*o*_ within the range of [**0**, 0.6***G***], we found that the prior model relying on prediction exhibits errors that are independent of ***σ***_*o*_; the likelihood model relying on observation exhibits errors that increase with ***σ***_*o*_; and the posterior model exhibits smaller errors than either the prior or the likelihood models, similar to the EKF agent (Fig. 7**b**; see Methods, agent selection). This suggests that the modular agent’s internal belief is formed by combining prediction and observation, weighted by their relative uncertainty, akin to the EKF. However, unlike the EKF that is provided with the ground truth values of uncertainties and the state transition, RNNs in the modular agent must learn to infer these from inputs (***o***_*t*_, ***a***_*t*−1_) in training.

### Relative reliability of information sources in training shapes generalization

Since an EKF-like belief update rule can emerge in modular agents’ RNNs after training, we explored how the bias in this rule favoring one information source over the other influences generalization. In all previous sections (except the last one), agents were trained with fixed ***σ***_*a*_ = 0.2***G, σ***_*o*_ = 0.1***G*** in the training task, learning to rely more on observation. Here, we trained 16 modular agents, each with a combination of ***σ***_*a*_ and ***σ***_*o*_ within {**0**, 0.1***G***, 0.2***G***, 0.3***G***} in the training task. We then tested their generalization performance (AUC) and corresponding belief accuracy (decoding error) in the novel gain and perturbation tasks (see Methods, agent selection). For both novel tasks, we found that agents trained with smaller observation uncertainties (***σ***_*o*_ *<* ***σ***_*a*_) generalized better than agents with equal uncertainties (***σ***_*o*_ = ***σ***_*a*_), and worst of all were agents trained with larger observation uncertainties (***σ***_*o*_ *>* ***σ***_*a*_; Fig. 7**c**–**d**). Decoding errors were larger for agents exhibiting poorer performance (Fig. S7**a**–**b**), evidence that poor generalization is associated with inaccurate internal beliefs (Fig. S7**c**–**d**).

This result is expected, given the structure of the novel tasks: agents must be aware of novel gains or perturbations via optic flow, since their internal model for prediction is outdated in novel tasks with novel state transitions. Agents trained with larger observation uncertainties tend to trust their prediction more and ignore changes that must be observed via visual cues in novel tasks (a mismatch between prior knowledge and the task structure). Therefore, like humans and macaques [23], agents must rely more on observation to generalize to these tasks.

## Discussion

The brain has evolved advantageous modular architectures for mastering daily tasks. Here, we investigated the impact of architectural inductive biases on learning and generalization using deep RL agents. We posited that employing a modular architecture with functionally specialized modules would allow agents to efficiently capture the structure of a given task during training, and then use this knowledge to support generalization in novel tasks with different parameters but a similar structure. To test this, we trained agents with different architectures on a partially observable navigation task and tested these agents with novel sensorimotor mappings or passive perturbations. We found that agents using a modular architecture not only learn the training task more efficiently, but their learned internal state representations of the environment remain accurate in novel tasks. Consequently, these agents exhibit superior generalization performance than those using architectures with less specialized modules.

We also found that modular agents learn a Kalman filter-like belief update rule, weighing the relative reliability of information sources for belief formation, similar to how the brain performs probabilistic inference [22]. Therefore, having higher uncertainty in observation than in prediction during training can bias agents toward dead-reckoning strategies, as they learn not to trust their observation. The agents are then less sensitive to novel patterns in their observation, and thus form wrong internal beliefs when tested in novel tasks, impeding their generalization abilities.

### Modular actor-critic RL models bridge neuroscience and AI

The orbitofrontal cortex (OFC) has been suggested to compute state representations by integrating sensory inputs in partially observable environments [31], then project these beliefs to the basal ganglia (BG) [32, 33]. One possible role for the BG is to construct the value function of a task using TD errors of the dopamine (DA) system [34], and then select the best action at each state proposed by the cortex that leads to the highest value [35, 36, 37, 38].

Inspired by these brain mechanisms, we developed an agent with modular critic and actor in a similar manner. The modular critic, analogous to the OFC-BG circuit, uses a dedicated module to compute belief states and passes these beliefs to a second module responsible for value computation. It is trained using the TD error. Meanwhile, the actor’s synaptic weights are optimized to generate actions that the critic evaluates favorably. Our results demonstrate that this modular critic enhances the agent’s understanding of the task structure, leading to improved performance in both training and novel tasks compared to agents without such specialized modules. This rationalizes the modularization of the brain’s OFC-BG circuit for RL. Furthermore, the advantages of modularization in actors justify the macaque brain’s utilization of multiple cortical areas (e.g., dorsomedial superior temporal area, parietal area 7a, dorsolateral prefrontal cortex [PFC]) to construct a control policy for this task [26] and achieve effective generalization [18].

Interestingly, our learning algorithm is only stable when the actor is updated more slowly than the critic (Methods; [29]), and this property could potentially explain why the BG learns from experience faster than the PFC [39]: if the critic’s value estimation hasn’t improved since the last training step, the actor/cortex cannot learn anything new from the critic/BG.

### Achieving generalization in RL

Different classes of RL models have different generalization abilities. The classic model-free RL algorithm, reminiscent of the brain’s DA system [34], selects actions that maximize the stored value function associated with the environmental model (state transition probabilities and reward functions) used in training. It does not adapt to any changes in the environmental model that should lead to a new value function. The successor representation algorithm [40, 41] decomposes the value into a representation of transition probabilities learned from experience and a reward function model, realizing immediate generalization on new rewards by reconstructing new values with learned transitions and new reward function models. However, upon encountering changes in transition probabilities, such as posed by our gain and perturbation tasks, the successor representation requires a similarly lengthy relearning as a model-free algorithm. On the other hand, the meta-RL algorithm [42] bypasses the tedious relearning of new value functions for novel tasks. Inspired by the standalone fast learning system of the PFC that is shaped by (but distinct from) the slow DA-based RL [43], meta-RL uses model-free values to train a standalone RNN policy network that maps inputs to actions across multiple tasks, considering current observations and previous actions and rewards as inputs. This allows the policy network to learn not just a single policy but an embedded learning algorithm. In novel tasks, the policy can explore different actions and gradually improve its performance by using reward feedback associated with those actions [43].

Macaques in our navigation task demonstrated instant generalization to task manipulations without relying on learning from novel trials [18]. This may be attributed to their inherent inductive biases evolved for natural tasks [26]. Similar to meta-RL, we used model-free values (critic) to train a standalone policy (actor). However, to emphasize instant generalization, we excluded the reward from the actor’s inputs. This prevents the actor from relying on reward feedback to improve its policy in new tasks. Nonetheless, with appropriate inductive biases, the actor can acquire an accurate understanding of the task structure and effectively map new observations from novel tasks to appropriate actions without further learning. Our modeling highlights the importance of harnessing inductive biases to facilitate efficient generalization in complex environments.

### Future directions

A key question connecting AI and neuroscience is how to reconcile the various distinct RL models that exhibit analogs in the brain and behavior. For example, it is believed that the midbrain DA neurons implement model-free RL through trial and error [34]. The successor representation has the ability to replicate place and grid cell properties [40]. The learning-to-learn system in PFC is explained by meta-RL [42]. The flexibility of humans and animals in planning suggests the presence of model-based RL [44]. Furthermore, the brain possesses unique inductive biases evolved for natural tasks, allowing animals to achieve instant adaptation, as explored by our model. The brain’s reward-based learning system could consist of a family of overlapping algorithms implemented by different brain regions, such that each of the aforementioned models captures a specific aspect of our cognitive capacities. Animal behavior may inherit properties from multiple algorithms [45] or selectively adopt the most appropriate algorithm based on the current task demands [46]. Future studies should aim to harmonize the diverse findings in RL literature in order to develop a more comprehensive understanding of the brain’s learning mechanisms.

It is also worth noting that the brain acquires module specialization through two distinct mechanisms. At the architectural level, module specialization is enforced as an inductive bias among functionally specialized regions [6]. At the representational level, even within the same region, functionally specialized modular clusters can emerge through learning [47]. While it is theoretically possible for homogeneous networks to internally develop functionally distinct clusters and achieve similar modularization as that enforced through architectural design, the performance difference for our agents with different architectures suggests that this phenomenon may not occur in our task. One possible explanation could be the nature of the task: specialized neural clusters are more likely to emerge when agents are exposed to multiple tasks with shared variables, rather than a single task [48]. Another possibility is that the standard optimization methods we used may be insufficient for networks to discover module-specialized solutions. One potential remedy is to introduce more realistic constraints during optimization. For example, embedding neurons in physical and topological spaces where longer connections incur higher costs could encourage clusters to form [49]. In the future, it would be valuable to investigate how to induce networks to learn modular clusters that are comparable to architectural modules. It would also be intriguing to explore the advantages of incorporating both types of modules. These directions hold the potential to deepen our understanding of the brain’s learning and generalization mechanisms and inspire the development of more efficient and adaptable AI systems.

## Methods

### Task structure

The navigation task and its manipulations were originally designed for macaques [16, 17, 18, 23, 26, 27]. All macaque data used in this paper were from previous works [16, 17], where details of the animal experiment setup can be found. We modeled this task as a POMDP [28] for RL agents, containing a state space *S*, an action space *A*, a transition probability *T*, a reward function *R*, an observation space Ω, an observation probability *O*, and a temporal discount factor *γ* = 0.97 over steps within a trial.

Each state ***s***_*t*_ ∈ *S* is a vector 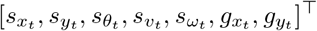 containing the agent’s x and y positions (cm), head direction (^*◦*^), linear and angular velocities (cm/s, ^*◦*^/s), and the target’s x and y positions (cm). The initial state of each trial was defined as 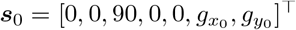, since the agent always starts from the origin facing forward (90^*◦*^). The target location was uniformly sampled on the ground plane before the agent, with the radius *g*_*r*_ ∈ [100 cm, 400 cm] and the angle *g*_*θ*_ ∈ [90^*◦*^ −35^*◦*^, 90^*◦*^ + 35^*◦*^] relative to the agent’s initial location. Specifically, angles were drawn uniformly within the field of view, *g*_*θ*_ ∼ 𝒰 (55^*◦*^, 125^*◦*^), and we sampled radial distances as 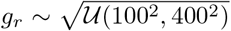 to ensure a spatially uniform distribution in 2D. Target positions in Cartesian coordinates are then 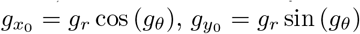.

Each action ***a***_*t*_ ∈ *A* is a vector 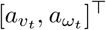 containing the agent’s linear and angular joystick actions, bounded in [−1, 1] for each component.

State transitions ***s***_*t*+1_ ∼ *T* (***s***_*t*+1_|***s***_*t*_, ***a***_*t*_) were defined as ***s***_*t*+1_ = ***f***_env_(***s***_*t*_, ***a***_*t*_) + ***η***_*t*_, where

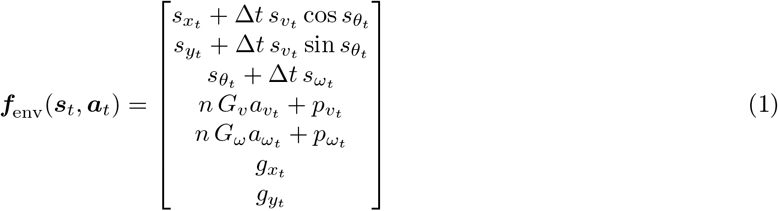

and zero-mean independent Gaussian process noise added to the velocities

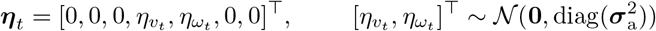

with standard deviation 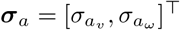 The operator diag(·) constructs a diagonal matrix with its vector argument on the diagonal. The timestep is ∆*t* = 0.1 s. Joystick gain ***G*** = [*G*_*v*_, *G*_*ω*_]^⊤^ = [200 cm/s, 90^*◦*^/s]^⊤^ maps dimensionless linear and angular joystick actions to units of velocities. Gain multiplier *n* scales ***G***. Linear and angular perturbation velocities are 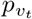and 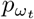.

The reward function *R*(***s***_*t*_, ***a***_*t*_) maps a state-action pair to a scalar *r*_*t*_. We firstly defined an action threshold *a*^∗^ = 0.1 to distinguish between when the agent had not yet begun moving, and when they moved and stopped: the agent must increase the magnitude of at least one action component above *a*^∗^ in the beginning (start criterion), then the agent must reduce the magnitude of both action components below *a*^∗^ to indicate a stop (stop criterion). Non-zero rewards were only offered in the last step of each trial and if the agent satisfied both criteria. For the non-zero rewards, we defined 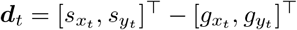 as the displacement between the agent’s and the target’s locations, and a reward *r*_*t*_ = 10 would be given if the Euclidean distance ||***d***_*t*_||_2_ was smaller than the radius of the reward zone *d*^∗^ = 65 cm. To facilitate training in the early stages, we allowed a small reward 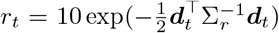 if the agent stopped outside the reward zone, where 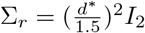 is a constant matrix, and *I*_2_ denotes the identity matrix of size 2.

A trial ended when the agent stopped, or if *t* exceeded the maximum trial duration 3.4 s. For later convenience, let *D*_*t*_ denote a trial completion flag that equals 1 if the trial is done at *t*, otherwise 0. A new trial thereafter started with a new sampled initial state ***s***_0_.

Observation ***o***_*t*_ ∈ Ω is a vector 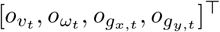 containing observations of the agent’s linear and angular velocities through optic flow, and the target’s x and y positions when visible in the first 0.3 s of each trial. ***o***_*t*_ ∼ *O*(***o***_*t*_|***s***_*t*_) was defined as

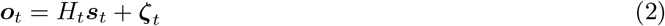

where ***ζ***_*t*_ is a zero-mean Gaussian observation noise, and the observation model *H*_*t*_ is a 4 × 7 matrix filled mostly with zeros, except for a few observable elements depending on the time within a trial: When *t* ≤ 0.3 s, the target is visible, so *H*^1,4^, *H*^2,5^, *H*^3,6^, *H*^4,7^ are equal to 1, where superscripts denote row and column; after *t* = 0.3 s the target disappears and only the optic flow is observable, so only *H*^1,4^, *H*^2,5^ are 1. For the observation noise, 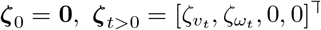, where 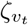and 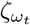 denote linear and angular observation noises, and 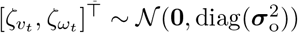 with standard deviation 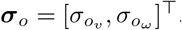.

### Task parameters

#### Training task

The gain multiplier is given by *n* = 1. There were no perturbations, so for any 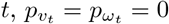. Process and observation noise standard deviations were in units of ***G***, i.e., ***σ***_*a*_ = *α*_*a*_***G, σ***_*o*_ = *α*_*o*_***G***, where *α*_*a*_ = 0.2, *α*_*o*_ = 0.1 were used to train agents before Fig. 7; *α*_*a*_, *α*_*o*_ ∈ [0, 0.3] were used in Fig. 7 except Fig. 7**b**.

#### Gain task

The gain multiplier *n* was 1.5 or 2 for analyses before Fig. 6, or was sampled in 𝒰(3, 4) for the remaining analyses. Noise standard deviations were also multiplied by the same gains, i.e., ***σ***_*a*_ = *α*_*a*_*n****G, σ***_*o*_ = *α*_*o*_*n****G***. There were no perturbations.

#### Perturbation task

Parameters *n*, ***σ***_*a*_, ***σ***_*o*_ were the same as those in the training task. There were three perturbation parameters uniformly sampled for each trial: perturbation peak time relative to the trial start *t*_*p*_ ∼ 𝒰 (0.5 s, 1.5 s), and perturbation linear and angular peaks 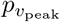and 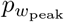. These peaks were sampled from the ranges specified in the caption for each analysis. The sampled perturbation parameters determined Gaussian-shaped linear and angular perturbations, defined as

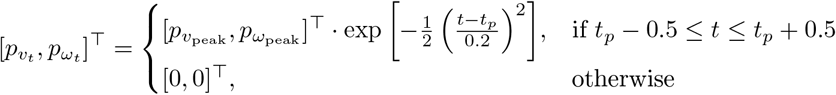

#### Uncertainty task

The gain multiplier was *n* = 1, and there were no perturbations. The prior model was trained with ***σ***_*a*_ = **0, *σ***_*o*_ = 0.8***G***; the likelihood model was trained with ***σ***_*a*_ = 0.8***G, σ***_*o*_ = **0**; the posterior and the EKF models were trained with ***σ***_*a*_ = 0.3***G*** and a randomly varying ***σ***_*o*_ = *α*_*o*_***G*** where *α*_*o*_ ∼ 𝒰 (0, 1) was drawn independently for each trial. A standard deviation of **0** denotes the noise-free case.

## Belief modeling

The state ***s***_*t*_ is partially observable in our task, therefore, an agent cannot decide on ***a***_*t*_ only based on the current sensory inputs. It can internally maintain a belief state representation *b*_*t*_, which is a posterior of ***s***_*t*_, for decision-making. We considered both a model-based inference method and a gradient-based optimization method to model the belief.

### Recursive Bayesian estimation

When the transition probability *T* and the observation probability *O* are known, the belief is a posterior of ***s***_*t*_ given all available observations and actions, i.e., *b*_*t*_ = *p*(***s***_*t*_ ***o***_0:*t*_, ***a***_0:*t*−1_), and can be inferred recursively as

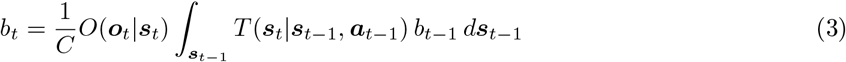

where *C* = *p*(***o***_*t*_|***o***_0:*t*−1_, ***a***_0:*t*−1_) is a normalization constant, and *b*_*t*−1_ = *p*(***s***_*t*−1_|***o***_0:*t*−1_, ***a***_0:*t*−2_).

#### EKF belief

When all variables are Gaussian in the Recursive Bayesian estimation and *T* is nonlinear, the EKF [30] is a tractable method that uses a local linearization to approximate eq. (3). The belief here is a Gaussian density *b*_*t*_ = *𝒩* (***ŝ***_*t*_, *P*_*t*_). To simplify the computation here, we express position in relative coordinates by letting the initial belief mean be 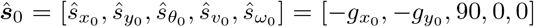, and let the state transition ***f*** _env_ contain the first five equations in eq. (1) to reduce the dimensionality of the state by two. Let *ϵ* denote a small number 10^−8^, we defined the initial belief covariance *P*_0_ = *ϵI*_5_. Let a 5 *×* 5 matrix *N*_*a*_ denote the Gaussian process noise covariance filled with 0 except 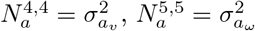. The observation’s dimensionality was reduced by two by omitting the target location, yielding 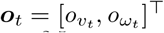. The observation model *H* in eq. (2) then becomes a 2 × 5 matrix filled with 0, except *H*^1,4^ = *H*^2,5^ = 1. Let 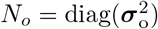denote the Gaussian observation noise covariance. Any 0 variance components in 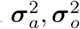 were replaced with a minimal variance of *ϵ* for *N*_*a*_, *N*_*o*_.

***f***_env_ at *b*_*t*−1_ = *𝒩* (***ŝ***_*t*−1_, *P*_*t*−1_) was locally linearized as

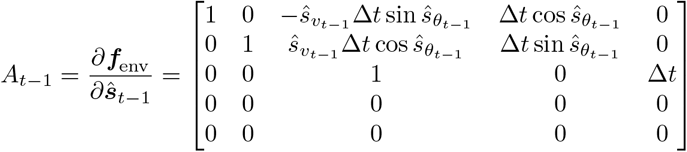

The EKF’s prediction step (eq. (4)) uses ***a***_*t*−1_ to get a predicted belief *b*_*t*|*t*−1_. Note that given our *A*_*t*−1_, velocity variance elements in the prediction 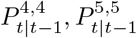 only depend on *N*_*a*_.

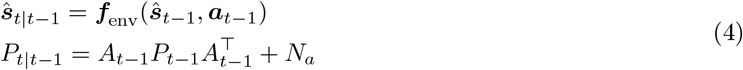

The EKF’s update step (eq. (5)) uses *b*_*t*|*t*−1_ and ***o***_*t*_ to get the final belief *b*_*t*_ = *𝒩* (***ŝ***_*t*_, *P*_*t*_). *K*_*t*_ is known as the Kalman gain which specifies the relative weights of prediction and observation. Mathematically, it is only affected by *N*_*a*_ and *N*_*o*_ and is independent of *P*_*t*−1_ in our task. As intuitively, past velocities do not predict current velocities except via the agent’s actions, which it knows.

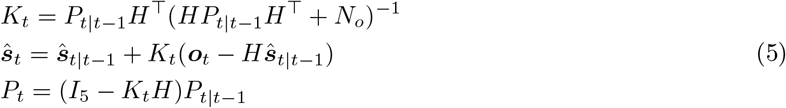

#### RNN belief

When the transition and the observation probabilities *T, O* are unknown to the agent, to support decision-making, an internal belief could be approximated via gradient-based optimization. We used RNNs to integrate partial observations ***o***_*t*_ and motor efference copies ***a***_*t*−1_ over time, trained end-to-end using the RL objective in our task (see below). It was shown in the result that an EKF-like belief update rule could emerge in RNNs, and therefore, the belief *b*_*t*_ resides in the RNN’s hidden state ***h***_*t*_. Each RNN maintains a hidden state ***h***_*t*_ = ***f*** _RNN_(***o***_*t*_, ***a***_*t*−1_, ***h***_*t*−1_) or ***h***_*t*_ = ***f*** _RNN_(***o***_*t*_, ***a***_*t*−1_, ***h***_*t*−1_, ***a***_*t*_) depending on its inputs (Fig. 6**a**–**b**). *b*_*t*_ encoded implicitly in ***h***_*t*_ is used by other neurons to compute ***a***_*t*_ or *Q*_*t*_ in the actor or critic.

### RL with EKF belief

Our RL algorithm for training the EKF agent with an EKF belief is based on an actor-critic approach called the Twin Delayed Deep Deterministic Policy Gradient (TD3) [29], referred to as EKF-TD3. We first computed beliefs using EKF as described above, and then trained neural networks to use those beliefs as inputs to guide actions.

#### Networks

Each agent has two critics with identical architectures but different initial weights to address the maximization bias in value estimation (see the critic update section below and [50]), although in the main text we only showed one of the critics used to train the actor to generate actions. Let ***i***_*t*_ denote the state-related inputs. All neural networks in an EKF agent were feed-forward, provided with the mean and covariance of *b*_*t*_ computed by the EKF, i.e., ***i***_*t*_ = {***ŝ***_*t*_, *P*_*t*_}. The actor and two critics are ***a***_*t*_ = ***π***_*μ*_(***i***_*t*_), 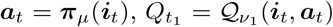, 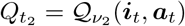, where *μ, ν*_1_, *ν*_2_ denote neural parameters.

#### Exploration

Since our actor is a deterministic function, to realize exploration in training, we combined the actor’s output with a zero-mean Gaussian exploration noise ***β***_*t*_, and clipped the sum to the box [−1, 1]:

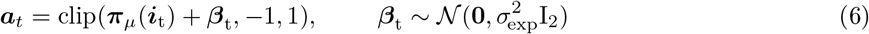

To ensure the agent can properly stop without noise variability, we let ***β***_*t*_ = **0** if the actor’s output ***π***_*μ*_(***i***_*t*_) is below the action threshold.

#### Experience replay

Instead of learning on the current trial, we used off-policy RL by storing experience in a replay buffer *ℬ*and frequently sampling data from *ℬ*to train the agent. At each state ***s***_*t*_, the EKF computed ***i***_*t*_ for the actor to generate ***a***_*t*_ following eq. (6). The agent observed the reward *r*_*t*_, next input ***i***_*t*+1_, and trial completion flag *D*_*t*_, and stored the one-step transition tuple (***i***_*t*_, ***a***_*t*_, *r*_*t*_, ***i***_*t*+1_, *D*_*t*_) in *ℬ*. The buffer *ℬ* had a capacity of 1.6*×* 10^6^ transitions, storing data on a first-in, first-out (FIFO) basis. Furthermore, we augmented the experience by also storing the mirror transition (*î*_*t*_, ***â***_*t*_, *r*_*t*_, *î*_*t*+1_, *D*_*t*_) generated by reflecting the original data across the y-axis.

#### Target networks

The learning of value in TD3 is akin to deep Q-learning [51]. Using the Bellman equation, ideally, the agent can learn to estimate the value 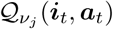by regressing the learning target 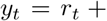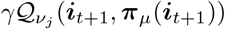, i.e., the one-step bootstrapping of the value after receiving the reward *r*_*t*_, observing the next input ***i***_*t*+1_, and estimating the next action ***π***_*μ*_(***i***_*t*+1_). One stability issue here is that the neural parameters for optimization are also used to construct the learning target *y*_*t*_, which changes at each learning step. To obtain a more stable *y*_*t*_, we thus maintained a copy of actor and critic networks with more slowly changing parameters *μ*^*′*^ and 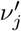 used in *y*_*t*_, referred to as target actor and critic networks. These parameters were initialized to be the same as *μ, ν*_*j*_ and passed through an exponential moving average,

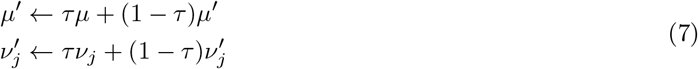

We used *τ* = 0.005.

#### Critic update

We sampled a batch of *M* = 256 transitions from the buffer each time,

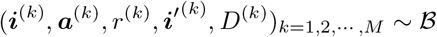

where the temporal subscript is omitted, and ***i***^*′*(*k*)^ denotes the next input after ***i***^(*k*)^. The next action given ***i***^*′*^(*k*) was estimated by the target actor network as

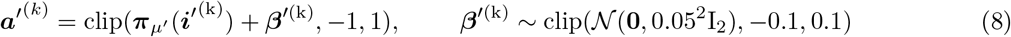

where ***β***^*′*(*k*)^ is small zero-mean Gaussian noise clipped to [−0.1, 0.1] to smooth the action estimation.

The learning target *y*^(*k*)^ used the smaller value estimation between two target critics to reduce the maximization bias [50], and was truncated at the end of each trial (*D*^(*k*)^ = 1). The learning objective of the two critics, *J*(*ν*_*j*_), *j* = 1, 2, was to regress the learning target *y*^(*k*)^, defined as

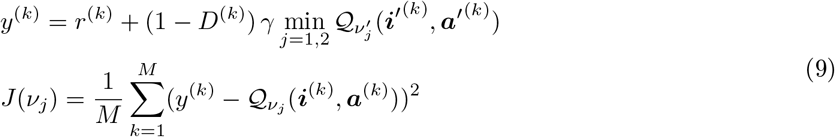

The gradient 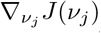 was computed by backpropagation (BP). Critic parameters *ν*_*j*_ were updated (see the agent training section below for optimizers) using 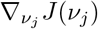 to minimize *J*(*ν*_*j*_).

#### Actor update

The actor’s parameter *μ* was updated once for every two critic updates. The actor’s learning objective *J*(*μ*) was to maximize the value of the first critic, defined as

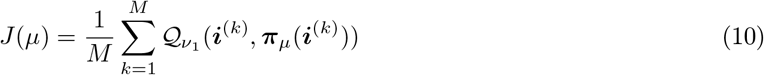

The gradient *∇* _*μ*_*J*(*μ*) was computed by BP. The actor parameter *μ* was updated using *∇*_*μ*_*J*(*μ*) to maximize *J*(*μ*). Note that the critic parameter *ν*_1_ was not updated here.

### RL with RNN belief

We developed a memory-based TD3 model leveraging RNNs to construct a form of internal beliefs to tackle POMDPs, referred to as RNN-TD3. All agents except the EKF agent were trained by this algorithm.

#### Networks

Let ***i***_*t*_ = {***o***_*t*_, ***a***_*t*−1_} and ***h***_*t*_ denote the state-related inputs and the RNN’s hidden state. The actor and two critics are 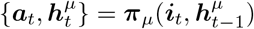, 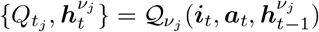, *j* = 1, 2, where we interpret that the belief *b*_*t*_ is implicitly encoded in all ***h***_*t*_ evolving over time. At the beginning of each trial, ***h***_*t*−1_ and ***a***_*t*−1_ were initialized to zeros. For simplicity, we drop ***h***_*t*_ in our notations for all networks’ outputs.

#### Exploration

Similar to that of EKF-TD3 (eq. (6)), we added zero-mean Gaussian exploration noise to the output of the actor during training if the output is above the action threshold,

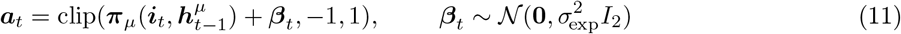

#### Experience replay

Similar to that of EKF-TD3, but rather than storing one-step transition tuples, the replay buffer *B* stored the whole trajectory for each trial of *N* time steps,

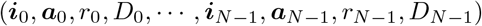

and its mirror image, because RNNs have hidden states ***h***_*t*_ generally depend on the entire history of inputs, not just the most recent ones. Each action was obtained using eq. (11). The FIFO buffer had a capacity of 10^5^ trajectories.

#### Target networks

Same as that of EKF-TD3.

#### Critic update

Similar to that of EKF-TD3, but critics here also needed to learn the temporal structure. Since the trial duration *N* varies across trials, we first sampled a trial duration 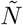 from the buffer ℬ, then sampled a batch of *M* = 16 trajectories with the same duration 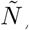,

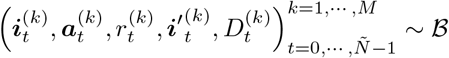

where 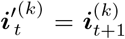. The next action 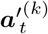, the learning target 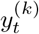, and the learning objective of the two critics *J*(*ν*_*j*_) were

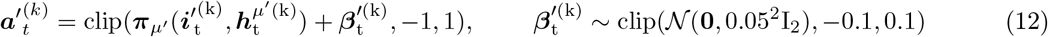

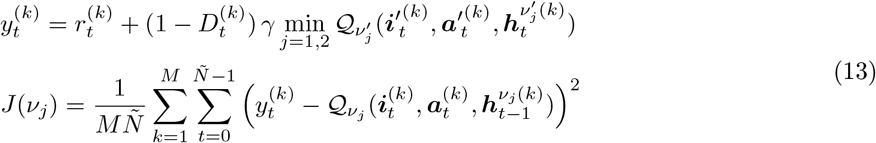

The gradient 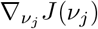 was computed by BP through time (BPTT). Critic parameters *ν*_*j*_ were updated using 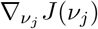 to minimize *J*(*ν*_*j*_).

#### Actor update

Similar to that of EKF-TD3, but the actor here needed to learn the temporal structure. The actor’s learning objective *J*(*μ*) was

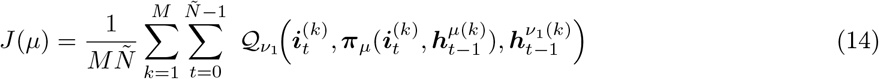

The gradient *∇*_*μ*_*J*(*μ*) was computed by BPTT. The actor parameters *μ* were updated using *∇*_*μ*_*J*(*μ*) to maximize *J*(*μ*).

### Agent training

All network parameters *μ, ν*_1_, *ν*_2_ were updated by the Adam optimizers [52]. Optimizer parameters were set as follows: learning rates annealed from 3*×* 10^−4^ to 5*×* 10^−5^, exponential decay rates for the first and second moment estimates = 0.9, 0.999, a constant added in denominators for numerical stability = 1.5 *×* 10^−4^, and weight decay = 0. The critics were updated once for every *c* = 4 interactions with the environment. The actor was updated once for every two critic updates.

During training, we periodically validated the agent’s performance with 300 validation trials, and used the moments when the agent achieved 20% and 80% accuracy to split the whole training course into three phases. The learning rates for the actor and critics, the exploration noise *σ*_exp_ (eq. (6),(11)), and the observation noise ***σ***_*o*_ (eq. (2)) in each phase were set as follows: In phase I, learning rates were 3*×* 10^−4^, *σ*_exp_ = 0.8, ***σ***_*o*_ = **0**. In phase II, learning rates were 3*×* 10^−4^, *σ*_exp_ = 0.5, ***σ***_*o*_ = *α*_*o*_***G***, where *α*_*o*_ is defined in the training or uncertainty tasks. In phase III, learning rates were 5 × 10^−5^, *σ*_exp_ = 0.4, ***σ***_*o*_ = *α*_*o*_***G***. Training was stopped after the agent had experienced 10^4^ trials after phase I for agents before Fig. 6, or 10^5^ trials for the remaining agents.

We summarize the EKF/RNN-TD3 algorithms for training all our agents in Algorithm 1.

#### Algorithm 1 EKF/RNN-TD3

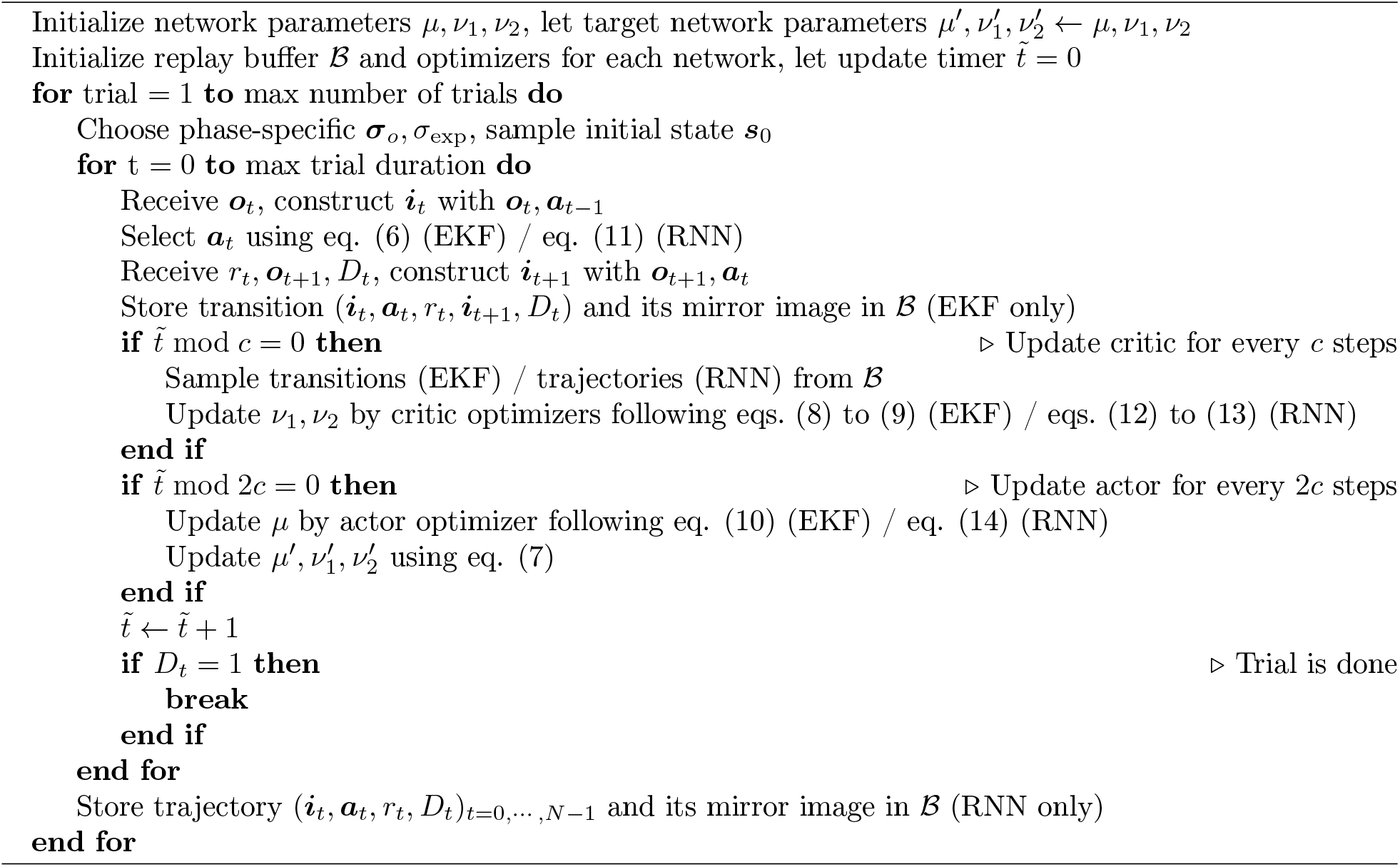

### Agent testing and TD errors

During testing, the target networks were no longer used. The trained actor ***π***_*μ*_ was used to interact with the environment and generate transition tuples (***i***_*t*_, ***a***_*t*_, *r*_*t*_, ***i***^*′*^_*t*_, *D*_*t*_) for each *t*. No exploration noise was added to the output of the actor.

The TD error shown in analyses is similar to the learning objective for critics (eq. (13)), except that the target networks were replaced with the trained networks, and only the first critic was used. Specifically, for each *t*, the next action is 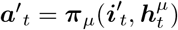, and the TD error is given by:

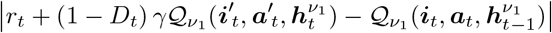

### Agent selection

During phases II and III training, every 500 training trials we saved neural parameters of each network. To fairly compare agents’ performance in each task (training, gain, perturbation, uncertainty), we tested all sets of stored parameters for each task with one or multiple test sets with 500 (for agents before Fig. 6) or 300 trials each, and endowed each agent with the neural parameters that allowed it to achieve the highest reward rate (number of rewarded trials per second averaged across test sets) for each task. The test sets used for each task are as follows. Training task: one test set with the training task’s parameters. Gain task: three or four test sets with the gain= 1*×*, 1.5*×*, 2 *×* (before Fig. 6) or 1*×*, 2*×*, 3*×*, 4*×* (Figs. 6 and 7). Perturbation task: two test sets for agents before Fig. 6, one without perturbation and the other with perturbation parameters sampled from the ranges shown in Fig. 5**a**. For agents in Figs. 6 and 7, one additional test set was included, with perturbation ranges identical to those in Fig. 6**d**. Uncertainty task: three test sets with ***σ***_*a*_ = 0.3***G*** and ***σ***_*o*_ = {**0**, 0.4***G***, 0.8***G***}.

### Agent architectures

Although all agents had two architecturally identical critics, we only showed one in the main text (Fig. 1**e**, Fig. 6**b**). All RNNs were implemented as Long Short-Term Memory (LSTM) networks [53]. All MLP layers linearly transformed inputs and then applied ReLU nonlinearities. The output of critics *Q*_*t*_ was produced by a linear unit without any nonlinearity; the linear and angular control outputs of the actors ***a***_*t*_ were bounded to [−1, 1] by hyperbolic tangent nonlinearities. In the holistic critic/actor (Fig. 6**a**–**b**), there were 220 LSTM units. In all other architectures in Fig. 6**a**–**b**, each RNN module had 128 LSTM units, and each MLP module contained two layers with 300 ReLU units in each. All architectures, as a result, had a similar number of parameters (Fig. S6**a**). The EKF agent’s actor and two critics used the same architecture consisting of an MLP module with two layers, each with 300 ReLU units.

### No-generalization hypothesis

For each gain trial with a novel gain *n****G*** for *n >* 1, or for each perturbation trial with novel non-zero perturbation velocities 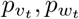, the hypothetical no-generalization trajectory was obtained as follows. We first recorded the agent/monkey’s sequential actions (***a***_0_, ***a***_1_, …, ***a***_*N*−1_) in the training task (1 *×* gain, no perturbations) navigating to the same target (for agents) or the closest target in the data set (for monkeys).

We then regenerated a new trajectory using (***a***_0_, ***a***_1_, …, ***a***_*N*−1_) following the environmental transition (eq. (1), process noise ***η***_*t*_ = 0), but with the novel gain multiplier *n* for the gain task or the novel perturbation velocities 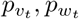 for the perturbation task.

### Under-/over-shooting definition using idealized circular trajectories

To determine when an agent or a monkey under- or over-shot the target in the gain task, we asked whether its stop location exceeded the target location in the distance along their corresponding idealized circular trajectories.

Specifically, given an arbitrary endpoint 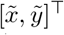, the circular trajectory connecting it from a forward heading (90^*°*^, initial head direction) at the origin (start location) has a radius as a function of this point 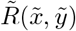. The arc length of this trajectory is a function:

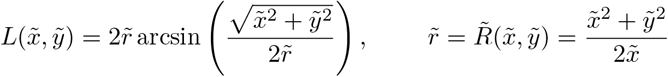

We deemed the agent’s stop location [*s*_*x N* −1_, *s*_*y N* −1_]^⊤^ to have overshot the target [*g*_*x*_, *g*_*y*_]^⊤^ if *L*(*s*_*x N* −1_, *s*_*y*−1_ *N*) *> L*(*g*_*x*_, *g*_*y*_), otherwise it undershot.

### Trajectory length and curvature

We approximated the length 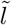 and the curvature 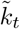of a trajectory 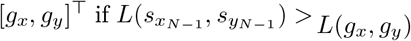 as follows:

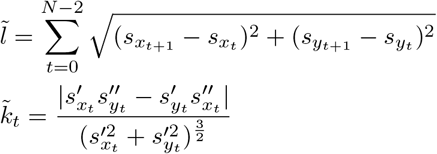

where first derivatives 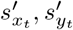 and second derivatives 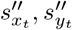were estimated using first-order one-sided differences for the first and last points and second-order central differences for interior points. In each trial, we excluded curvatures that surpassed the 95th percentile at any step, considering them as outlier values. Note that the monkeys’ trajectories here were downsampled to have the same 0.1 s time step as the agents’ trajectories.

### Spatial tuning

We obtained the approximate spatial tuning of each neuron by linearly interpolating its activity and the agent’s x and y location using data from each step across trials, followed by a convolution over the 2D space using a boxcar filter with height and width of 40 cm.

### Neural decoding

While agents were being tested, we recorded their sensory, latent, and motor variables for the analyses in Figs. 1**l** and S1**d** and their positions 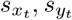 for all other decoding analyses. We also recorded their neural activities in each module for both their actors and critics. Let *S* denote a partitioned matrix where rows are time steps, and columns are decoding target variables, e.g., [***s***_*x*_, ***s***_*y*_] for agent’s positions. Recorded neural activities *X* were concatenated over time, where rows are time steps and columns are units. A linear decoder regressed *S* on *X*, whose partitioned parameters for all decoding variables *W* were obtained by the ridge estimator following

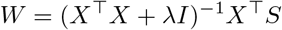

where *λ* is a penalty term chosen from {0.1, 1, 10} by cross-validation. Importantly, we always used 70% trials in the dataset to train the decoder, and used the remaining 30% trials to test the decoder’s predictions.

The decoding error of the belief in each trial was defined as

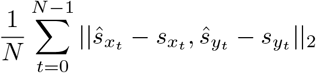

where 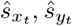are predicted x and y positions.

### Statistics

All agents were trained with 8 different random seeds, which determined the initialized neural network parameters and random variables in training (e.g., process and observation noises, agent’s initial state, exploration noise, and sampling from the buffer). All analyses for agents included data from training runs with all random seeds unless otherwise noted. We reported mean, SD, or SEM throughout the paper. All correlations were quantified by Pearson’s *r*.

In all violin plots, we determined upper and lower whiskers following *q*_1_ − whis (*q*_3_ − *q*_1_) and *q*_1_ +whis (*q*_3_ *– q*_1_), where *q*_1_, *q*_3_ are the first and third quartiles, and whis = 1.5 [54]. We did not plot outliers beyond the whisker range for better visualization, but we did not exclude them in quantification.

## Author contributions

Conceptualization and methodology: R.Z., X.P., and D.E.A.; formal modeling and analysis: R.Z.; original draft: R.Z.; review and editing: R.Z., X.P., and D.E.A.; funding acquisition: X.P. and D.E.A.; supervision: X.P. and D.E.A.

## Acknowledgements

We express our greatest appreciation to Kaushik Lakshminarasimhan for useful discussions. This work was supported by National Institutes of Health grants U19 NS118246 and R01 NS120407.

## Code availability

All training and analysis codes are available on GitHub https://github.com/ryzhang1/Inductive_bias.

## Data availability

All data used in this work are available from the corresponding author upon request.

## Supplementary figures

**Figure S1:**
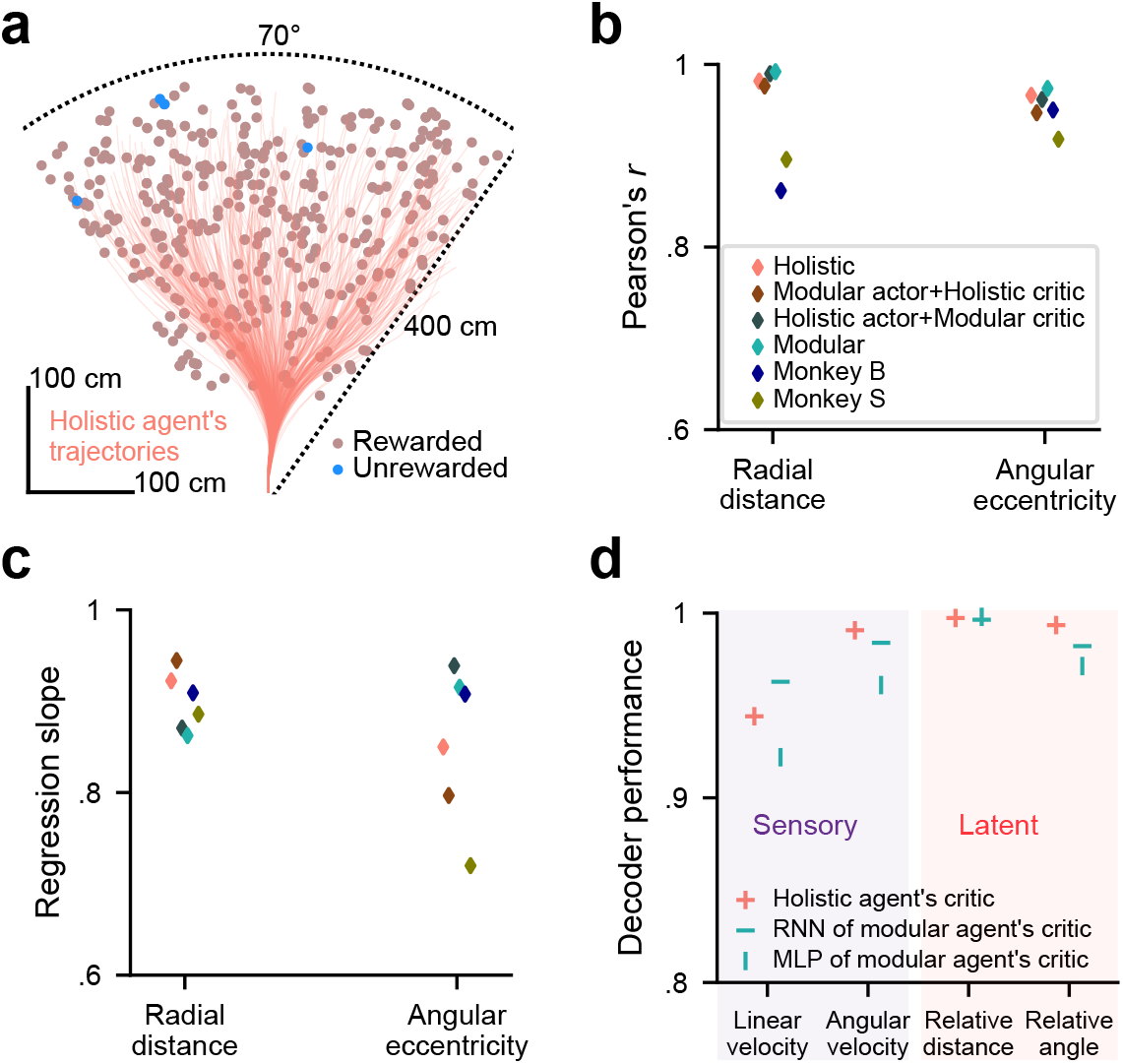
[Related to Fig. 1]. **a**. Similar to Fig. 1**h**, but showing an example holistic agent’s trajectories. **b**. Pearson correlation coefficient for agents’ and monkeys’ stop locations versus target locations after training, for data shown in Fig. 1**i. c**. Similar to **b**, but showing regression slopes (*>* 1/*<* 1: over-/under-shooting). **d**. Similar to Fig. 1**l**, but for modules in critics.

**Figure S2:**
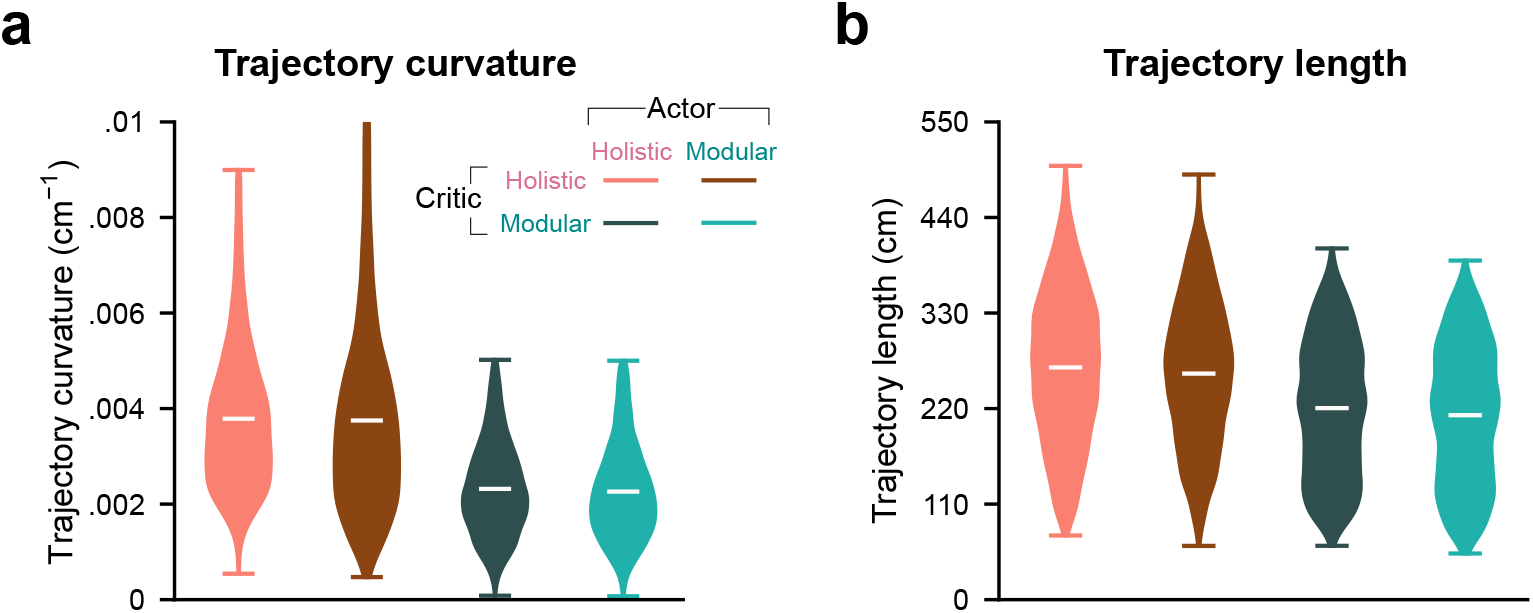
[Related to Fig. 2]. **a**. Distribution of trajectory curvature for agents navigating to the same set of 1000 targets. The curvatures were averaged across timesteps for each trajectory. **b**. Similar to **a**, but showing the length for each trajectory. **a**–**b**. Containing data from *n* = 8 random seeds for each agent. White bars denote means across trials.

**Figure S3:**
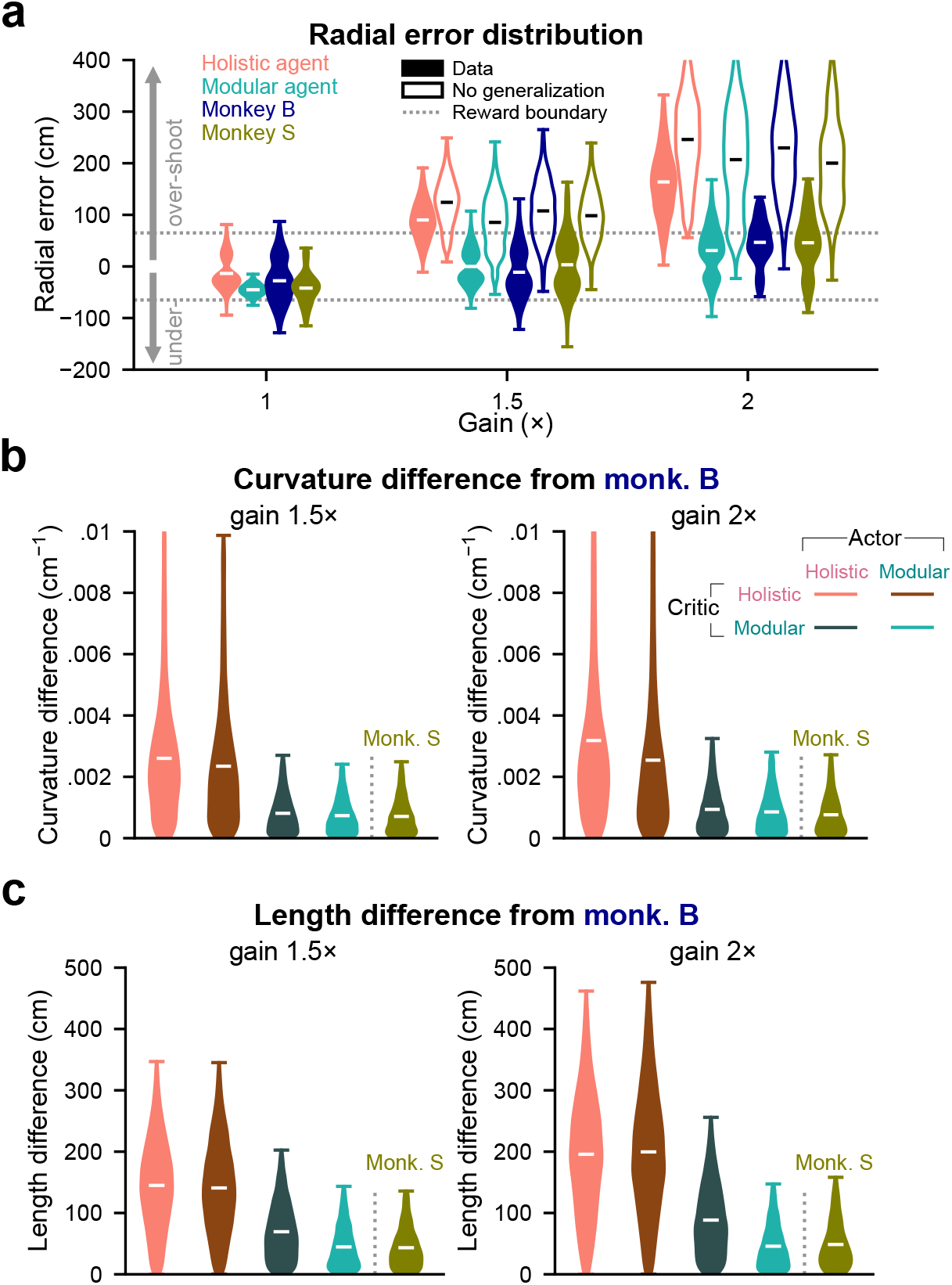
[Related to Fig. 3]. **a**. Similar to Fig. 3**c**–**d**, but showing detailed distributions of radial errors for agents and monkeys. **b**–**c**. Similar to Fig. 2**a**–**b**, but were conducted under two novel gain conditions. **a**–**c**. Containing data from *n* = 8 random seeds for each agent. White bars denote means across trials.

**Figure S4:**
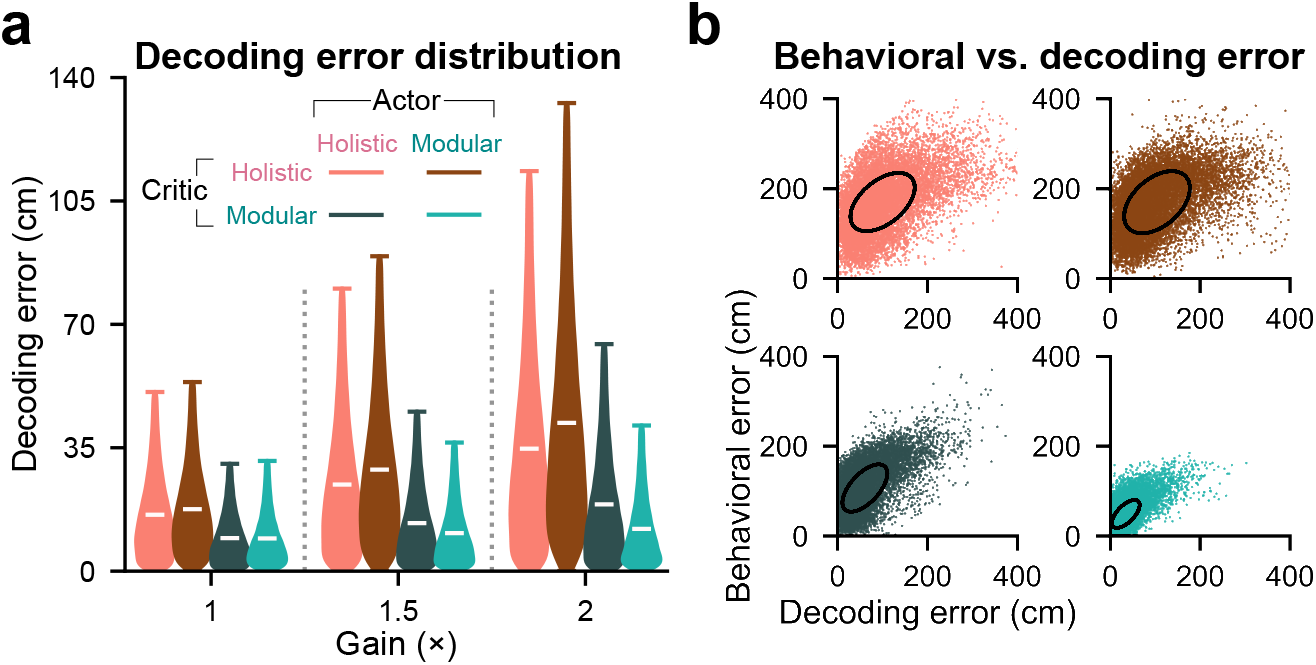
[Related to Fig. 4]. **a**. Similar to Fig. 4**d**, but showing detailed distributions of decoding errors. White bars denote means across trials. **b**. Behavioral error (absolute radial error between the target and the agent’s stop locations) versus decoding error of stop locations (distance between decoded and true stop locations) for trials used in Fig. 4**e**. Confidence ellipses capture 1 SD. Pearson’s *r*: Holistic, 0.50, Modular actor+Holistic critic, 0.45, Holistic actor+Modular critic, 0.61, Modular, 0.64, *p* = 0 for all agents. Regression slopes (without intercept): 1.28, 1.24, 1.31, 1.08. **a**–**b**. Containing data from *n* = 8 random seeds for each agent.

**Figure S5:**
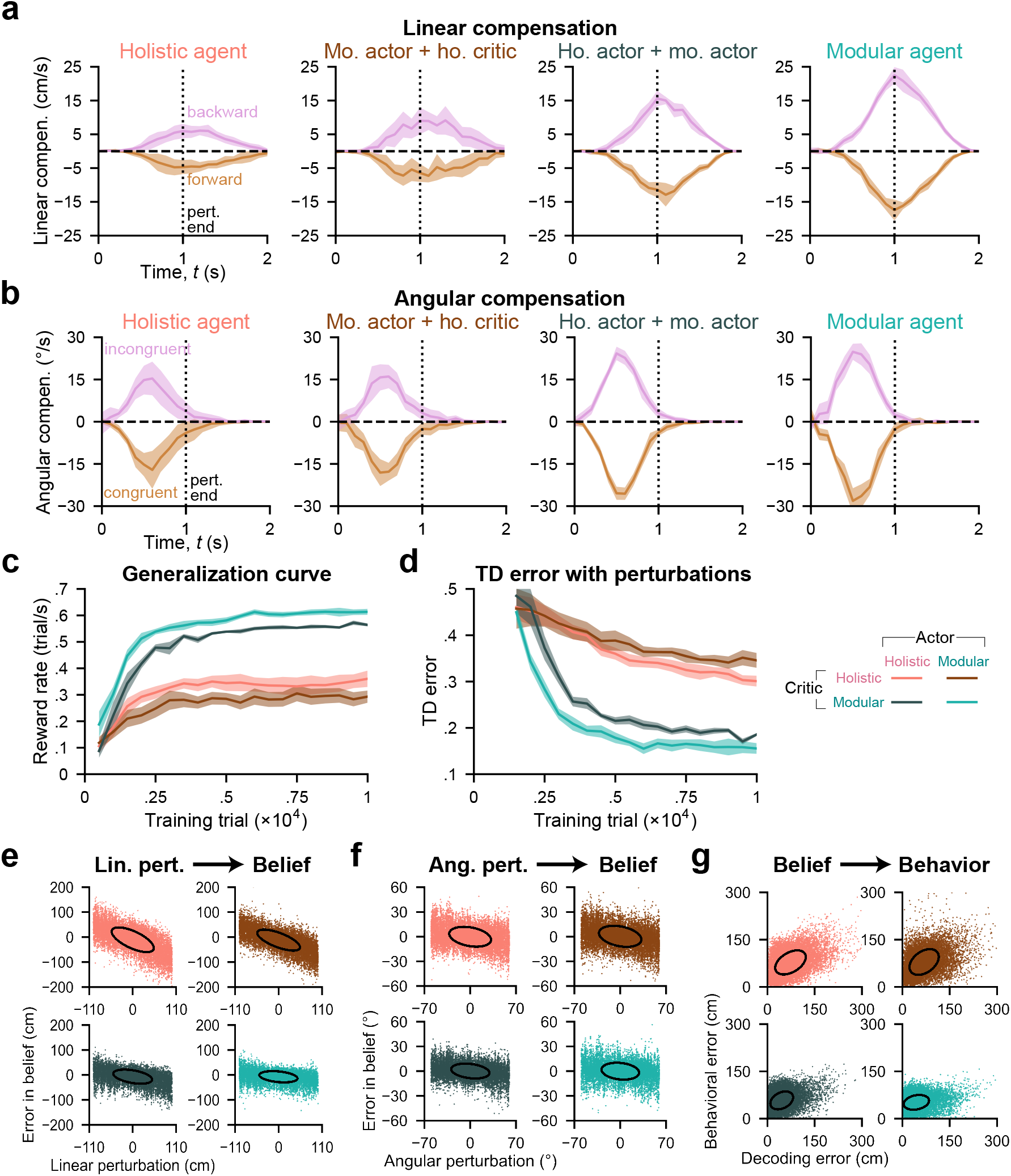
[Related to Fig. 5]. **a**–**b**. Agents’ linear (**a**) and angular (**b**) compensatory actions (in units of velocities) in a 2 s time window, averaged across a test set comprising 1000 trials with consistent random target locations and perturbation parameters for each agent. Trials are aligned such that perturbations start at *t* = 0 s. The linear/angular compensatory actions are obtained by subtracting linear/angular actions in target-matched unperturbed trials from those in corresponding perturbation trials. Perturbations causing one to get closer to/further from targets are grouped as forward (+)/backward (−) for linear perturbations (**a**) and congruent (+)/incongruent (−) for angular perturbations (**b**). Vertical dotted lines denote the perturbation end time (*t* = 1 s). Horizontal dashed lines denote the null compensatory action. Shaded regions denote *±*1 SD across *n* = 8 random seeds. **c**–**d**. Similar to Fig. 2**c**–**d**, but using a perturbation validation set. **e**–**f**. Same as the top row of Fig. 5**g**, but showing all data points. Each dot denotes a trial. **g**. Similar to Fig. S4**b**, but using trials in Fig. 5**h**. Pearson’s *r*: 0.45, 0.40, 0.34, 0.23, *p* = 0. Regression slopes (without intercept): 0.85, 0.90, 0.89, 0.70.

**Figure S6:**
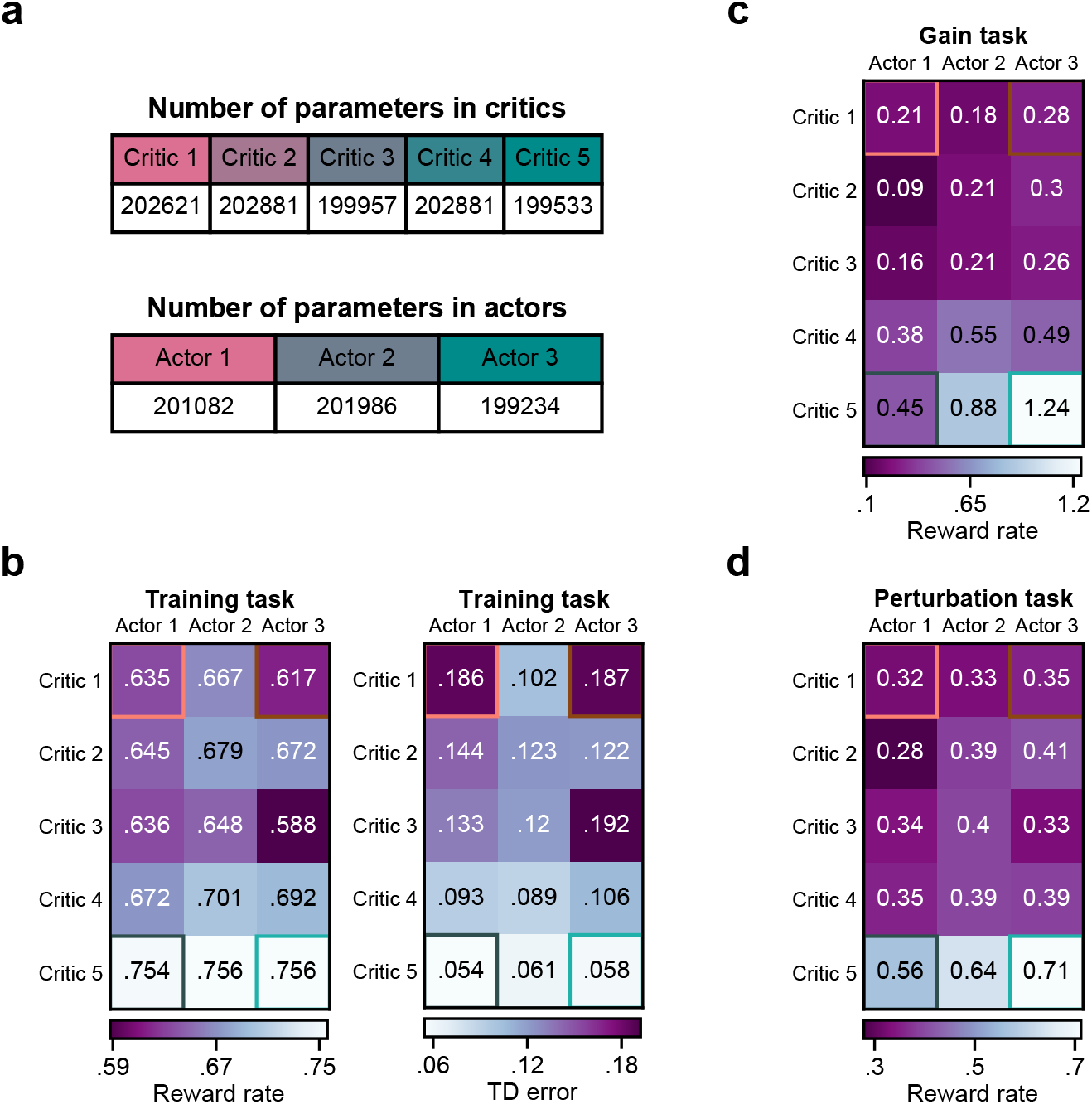
[Related to Fig. 6]. **a**. Total number of neural parameters for critics and actors in Fig. 6**a**–**b**. **b**. *Left*: Reward rate (rewarded trials/s) in 2000 training trials for agents with all combinations of actors and critics after training, averaged across *n* = 8 random seeds for each agent. *Right*: TD error (averaged cross steps and trials) evaluated using the trials on the left. The four corners represent the four agents used in the previous analyses. Text in white/black denotes that the agent is worse/better than the average value of all agents. Pearson’s *r* between left and right: − 0.93, *p* = 5.5 · 10^−7^. **c**–**d**. Similar to **b** left, but **c** uses gain trials from Fig. 6**c**, and **d** uses perturbation trials from Fig. 6**d**.

**Figure S7:**
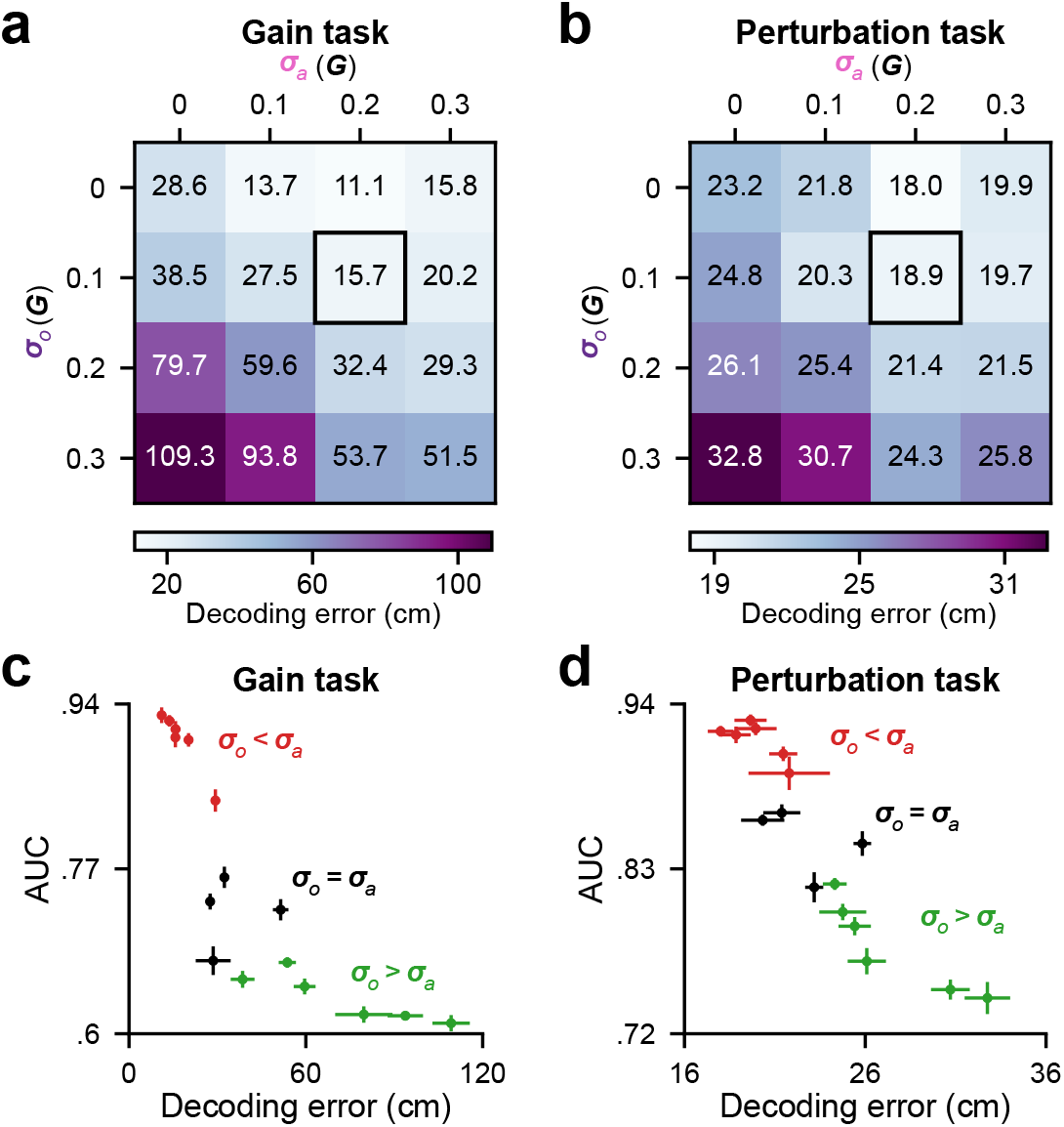
[Related to Fig. 7]. **a**–**b**. Similar to Fig. 7**c**–**d**, but showing the decoding error, averaged across time steps and trials. 70% trials were used for training the decoder, while the remaining trials were used for analyses. **c**–**d**. AUC (Fig. 7**c**–**d**) versus decoding error (**a**–**b**) for agents trained with all combinations of ***σ***_*a*_ and ***σ***_*o*_, tested in the gain (**c**) and the perturbation (**d**) tasks. Error bars denote *±* 1 SEM across *n* = 8 random seeds.

